# Screening of Tau Protein Kinase Inhibitors in a Tauopathy-relevant cell-based model of Tau Hyperphosphorylation and Oligomerization

**DOI:** 10.1101/821389

**Authors:** Hamad Yadikar, Isabel Torres, Gabrielle Aiello, Milin Kurup, Zhihui Yang, Fan Lin, Firas Kobeissy, Richard Yost, Kevin K. Wang

## Abstract

Tauopathies are a class of neurodegenerative disorders characterized by abnormal deposition of post-translationally modified tau protein in the human brain. Tauopathies are associated with Alzheimer’s disease (AD), chronic traumatic encephalopathy (CTE), and other diseases. Hyperphosphorylation increases tau tendency to aggregate and forms neurofibrillary tangles (NFT), a pathological hallmark of AD. In this study, okadaic acid (OA, 100 nM), a protein phosphatase 1/2A inhibitor, was treated for 24h in mouse neuroblastoma (N2a) and differentiated rat primary neuronal cortical cell cultures (CTX) to induce tau-hyperphosphorylation and oligomerization as a cell-based tauopathy model. Following the treatments, the effectiveness of different kinase inhibitors was assessed using the tauopathy-relevant tau antibodies through tau-immunoblotting, including the sites: pSer202/pThr205 (AT8), pThr181 (AT270), pSer202 (CP13), pSer396/pSer404 (PHF-1), and pThr231 (RZ3). OA-treated samples induced tau phosphorylation and oligomerization at all tested epitopes, forming a monomeric band (46-67 kDa) and oligomeric bands (170 kDa and 240 kDa). We found that TBB (a casein kinase II inhibitor), AR and LiCl (GSK-3 inhibitors), cyclosporin A (calcineurin inhibitor), and Saracatinib (Fyn kinase inhibitor) caused robust inhibition of OA-induced monomeric and oligomeric p-tau in both N2a and CTX culture. Additionally, a cyclin-dependent kinase 5 inhibitor (Roscovitine) and a calcium chelator (EGTA) showed conflicting results between the two neuronal cultures.This study provides a comprehensive view of potential drug candidates (TBB, CsA, AR, and Saracatinib), and their efficacy against tau hyperphosphorylation and oligomerization processes. These findings warrant further experimentation, possibly including animal models of tauopathies, which may provide a putative Neurotherapy for AD, CTE, and other forms of tauopathy-induced neurodegenerative diseases.

## Background

Tauopathy is a class of neurodegenerative condition that is associated with pathological phosphorylated tau protein accumulation in the human brain. Tauopathy has been associated with several clinicopathological conditions, including chronic traumatic encephalopathy (CTE) (1), traumatic brain injuries (2), post-traumatic stress disorder(3), and Alzheimer’s disease (AD)(4, 5).

Tau is a structural protein whose function is to promote microtubule stabilization and assembly, which are controlled by its phosphorylation state (6–8). In humans, the tau gene encodes the tau protein and is located on chromosome 17q21 (9). The main tau protein is encoded by 11 exons which are subjected to alternative splicing on exon two, three, and ten forming six isoforms. The six tau isoforms range from 352 to 441 amino acids. Tau isoforms vary in either having zero, one, or two N-terminal inserts (exons 2 and 3) and three or four repeats region at the C-terminal region (exon 10) (10, 11).

Tau protein consists of 79 potential phosphorylatable Serine and Threonine sites on the longest isoform. At least thirty tau phosphorylation sites have been reported in healthy conditions. Tau’s phosphorylation state and its ability to interact with microtubule proteins are regulated by various protein kinases and phosphatases (12, 13). Imbalances in the activities of tau kinases and phosphatases can cause tau to become hyperphosphorylated at specific residues leading to a higher tendency to dissociate from microtubules. Abnormally dissociated tau have a higher susceptibility of forming larger protein aggregates, filament assembly, and bundling of pair helical filaments (PHF) into neurofibrillary tangles (NFT) leading to cellular neurotoxicity(6–8, 14, 15).

Tau phosphorylation is carried out by a host of different kinases under physiological condition. Abnormal activities of tau kinases have been associated with AD, including kinases such as Src family kinase, Ca^2+^/calmodulin-dependent protein kinase II (CaMKII); cyclin-dependent kinase 5 (CDK5); casein kinase (1α/1δ/1ɛ/2); dual-specificity tyrosine kinase phosphorylation and regulated kinase-1A/2 (DYRK1A/2), glycogen synthase-3, and Fyn kinase(16, 17). Notably, a study reported that hippocampus and temporal cortex regions of the brain have high levels of CKII in AD when compared to controls (18). Furthermore, the tyrosine kinase Fyn has been highly researched for its implications with tau and neurodegeneration in the post-synapse N-methyl-D-aspartate receptors (NMDAR)(16, 19–21). Fyn phosphorylates tau in the N-terminal domain in neurons and plays a fundamental role in the amyloid signal transduction(16).

Several approaches for the treatment of tauopathic conditions have been investigated, including targeting tau kinases(22), activation of tau phosphatases(23), enhancing microtubule stabilization(24), tau immunotherapy(25), tau clearance(2), tau aggregation inhibition (26). Since *in vivo* tau-hyperphosphorylation results from multiple kinase activities, a single effective strategy to reverse tauopathies is still an open question. The inhibition of tau kinases using pharmaceutical drugs can lead to decreased levels of the hyperphosphorylated tau protein, thereby less aggregated tau (27–32). Several tau kinase inhibitors are in clinical trials for the treatment of tauopathies-related diseases (33). The most progressive protein kinase inhibition approach in the clinic thus far has been targeted at GSK-3β protein (30, 34).

It has been shown in AD and various other tauopathies that, tau is abnormally phosphorylated at Ser202, Ser396/404, Thr181, Thr205, and Thr231 (35, 36). The phosphorylation profile of tau residues at Ser202/Thr205 has been well-characterized in AD cases based on using specific antibodies (37). Analyzing these phosphorylation sites helps to show a pattern of relationships between tau protein phosphorylation and pathology.

Okadaic acid (OA), a protein phosphatase 1 and 2A (PP1/PP2A) inhibitor, induces tau hyperphosphorylation at pathological sites in both animal and cell-based models (38–40). OA inhibition of tau phosphatases allows the activation of multiple tau kinases, leading to its hyperphosphorylation (41, 42). Moreover, it has been shown that OA treatment in wild-type mice causes tauopathy-related abnormality in different regions of the brain (43).

In this study, mouse neuroblastoma culture (N2a) and rat primary cerebrocortical neuronal (CTX) culture was treated with OA, to induce tau hyperphosphorylation and oligomerization mimicking a tauopathy-relevant condition. In these experiments, we used the OA-induced tauopathy culture model to screen for different tau kinase inhibitors using immunoblotting and phospho-specific tau antibodies. Thus, it was hypothesized that using OA-induced tau hyperphosphorylation and aggregation as a tauopathy model to screen for kinase inhibitors would translate into putative neurotherapeutic targets for tauopathies-related disorders. Data from this work has shown that OA-induced tau hyperphosphorylation and oligomerization were inhibited by the different treatments. This side-by-side overview both highlights targets not well described, as well corroborates with data from targets previously studied, to be assessed in different relevant tauopathy-related *in vivo* models.

## Methods

### Phosphorylation Inhibitors

Ethylene glycol-bis(β-aminoethyl ether)-N,N,N’,N’-tetraacetic acid (**EGTA**) (Sigma-Aldrich, St-Louis, MO, USA), Dithiothreitol (**DTT**) (Sigma-Aldrich), Lithium chloride (**LiCl**) (Sigma-Aldrich), N-(4-methoxybenzyl)-N’-(5-nitro-1,3-thiazol-2-yl)urea (**AR-A014418**) (Sigma-Aldrich), (9S,10R,12R)-2,3,9,10,11,12-Hexahydro-10-hydroxy-9-methyl-1-oxo-9,12-epoxy-1H-diindolo[1,2,3-fg:3’,2’,1’-kl]pyrrolo[3,4-i][1,6]benzodiazocine-10-carboxylic acid methyl ester (**K252a**) (Sigma-Aldrich), (2R)-2-1-butanol (**Roscovitine**) (Sigma-Aldrich), 4,5,6,7-Tetrabromo-2-azabenzimidazole (**TBB**) (Sigma-Aldrich), 1-(7-methoxyquinolin-4-yl)-3-(6-(trifluoromethyl)pyridin-2-yl)urea (**A-1070722**) (Sigma-Aldrich), cyclosporine A (Sigma-Aldrich), N-(5-chloro-1,3-benzodioxol-4-yl)-7-[2-(4-methylpiperazin-1-yl)ethoxy]-5- (tetrahydro-2H-pyran-4-yloxy)quinazolin-4-amine (**Saracatinib**) (Selleck Chemicals, Houston TX), (5S,6R,7R,9R)-6-methoxy-5-methyl-7-(methylamino)-6,7,8,9,15,16-hexahydro-17-oxa-4b,9a,15-triaza-5,9-methanodibenzo[b,h]cyclonona[jkl]cyclopenta[e]-as-indacen-14(5h)-one (**STS**) (ab120056; Abcam, Cambridge, MA, USA), Z-Asp-2,6-Dichlorobenzoyloxymethyl Ketone (**Z-DCB**) (Cayman Chemical, Ann Arbor Michigan) and okadaic acid (Cell Signaling Technology, Danvers, MA). SNJ-1945 was a gift from (Senju Pharmaceutical Co. Ltd., Kobe, Japan) (Table 1).

**Table 1.**
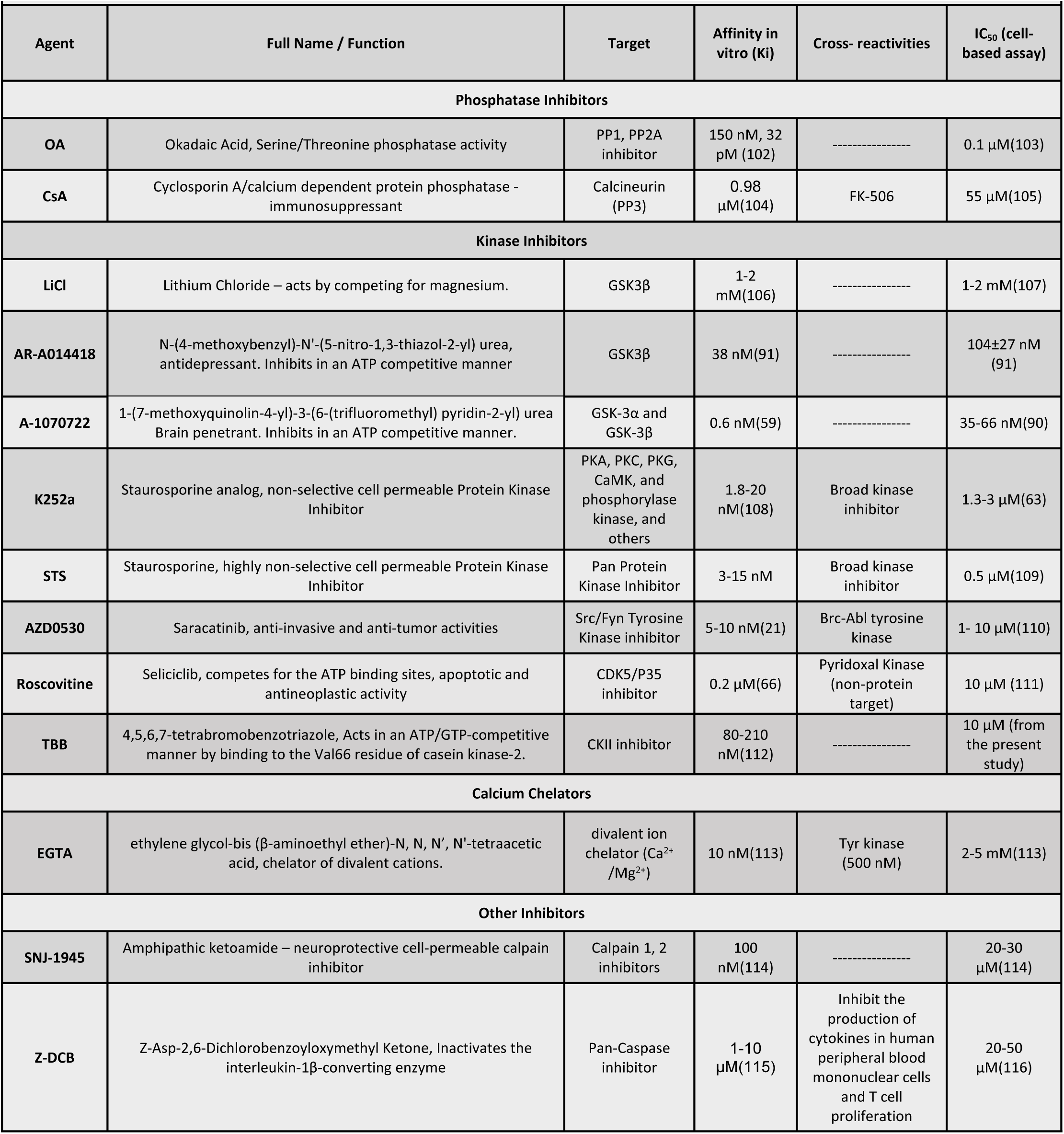
Phosphatase, Kinase inhibitor and other pharmacological agents used in the study.

**hyperphosphorylation in rat primary cerebrocortical neuronal cells.**
|  |  |  |  |  |  |
| --- | --- | --- | --- | --- | --- |
| Agent | Full Name / Function | Target | Affinity in vitro (Ki) | Cross- reactivities | IC<sub>50</sub> (cell-based assay) |
| Phosphatase Inhibitors |  |  |  |  |  |
| OA | Okadaic Acid, Serine/Threonine phosphatase activity | PP1, PP2A inhibitor | 150 nM, 32 pM (102) | ----- | 0.1 μM(103) |
| CsA | Cyclosporin A/calcium dependent protein phosphatase - immunosuppressant | Calcineurin (PP3) | 0.98 μM(104) | FK-506 | 55 μM(105) |
| Kinase Inhibitors |  |  |  |  |  |
| LiCl | Lithium Chloride – acts by competing for magnesium. | GSK3β | 1-2 mM(106) | ----- | 1-2 mM(107) |
| AR-A014418 | N-(4-methoxybenzyl)-N'-(5-nitro-1,3-thiazol-2-yl) urea, antidepressant. Inhibits in an ATP competitive manner | GSK3β | 38 nM(91) | ----- | 104±27 nM (91) |
| A-1070722 | 1-(7-methoxyquinolin-4-yl)-3-(6-(trifluoromethyl) pyridin-2-yl) urea Brain penetrant. Inhibits in an ATP competitive manner. | GSK-3α and GSK-3β | 0.6 nM(59) | ----- | 35-66 nM(90) |
| K252a | Staurosporine analog, non-selective cell permeable Protein Kinase Inhibitor | PKA, PKC, PKG, CaMK, and phosphorylase kinase, and others | 1.8-20 nM(108) | Broad kinase inhibitor | 1.3-3 μM(63) |
| STS | Staurosporine, highly non-selective cell permeable Protein Kinase Inhibitor | Pan Protein Kinase Inhibitor | 3-15 nM | Broad kinase inhibitor | 0.5 μM(109) |
| AZD0530 | Saracatinib, anti-invasive and anti-tumor activities | Src/Fyn Tyrosine Kinase inhibitor | 5-10 nM(21) | Bcr-Abl tyrosine kinase | 1- 10 μM(110) |
| Roscovitine | Selaciclib, competes for the ATP binding sites, apoptotic and antineoplastic activity | CDK5/P35 inhibitor | 0.2 μM(66) | Pyridoxal Kinase (non-protein target) | 10 μM (111) |
| TBB | 4,5,6,7-tetrabromobenzotriazole, Acts in an ATP/GTP-competitive manner by binding to the Val66 residue of casein kinase-2. | CKII inhibitor | 80-210 nM(112) | ----- | 10 μM (from the present study) |
| Calcium Chelators |  |  |  |  |  |
| EGTA | ethylene glycol-bis (β-aminoethyl ether)-N, N, N', N'-tetraacetic acid, chelator of divalent cations. | divalent ion chelator (Ca<sup>2+</sup> /Mg<sup>2+</sup>) | 10 nM(113) | Tyr kinase (500 nM) | 2-5 mM(113) |
| Other Inhibitors |  |  |  |  |  |
| SNJ-1945 | Amphipathic ketoamide – neuroprotective cell-permeable calpain inhibitor | Calpain 1, 2 inhibitors | 100 nM(114) | ----- | 20-30 μM(114) |
| Z-DCB | Z-Asp-2,6-Dichlorobenzoyloxymethyl Ketone, Inactivates the interleukin-1β-converting enzyme | Pan-Caspase inhibitor | 1-10 μM(115) | Inhibit the production of cytokines in human peripheral blood mononuclear cells and T cell proliferation | 20-50 μM(116) |

### Antibodies

The antibodies that were used in this study were: phospho-tau monoclonal antibodies PHF-1 (pSer396/pSer404, 1/1000), CP13 (pSer202, 1/1000), RZ3 (pThr231, 1/1000), AT8 (pSer202/pThr205, 1/1000), AT270 (pThr181, 1/1000) and total tau monoclonal antibodies: DA9 (a.a. 102-140, 1/1000), DA31 (aa150-190, 1/1000) (gift from Peter Davies, Albert Einstein College of Medicine, Bronx, NY), polyclonal total tau DAKO (aa243-441, 1/5000) (CiteAb, England). Mouse anti-αII-spectrin (ENZO Life Sciences, Farmingdale, NY, USA, 1/5000). β-actin was used as protein loading evenness control (abcam, Cambridge, MA, USA, 1/3000) (Table 2).

**Table 2.** Antibodies used in this study.

| Table 2. Antibodies used in this study |  |  |  |
| --- | --- | --- | --- |
| Clone name | Epitope* | Supplier (Catalog#) | MAb/ PAb |
| Phospho Tau antibodies |  |  |  |
| AT8 | pSer202/pThr205 | Fisher-Thermo (MN1020) | Mouse MAb |
| AT270 | pThr181 | Fisher-Thermo (MN1050) | Mouse MAb |
| RZ3 | pThr231 | Peter Davies, Albert Einstein College of Medicine, Bronx, NY | Mouse Mab |
| CP13 | pSer202 | Peter Davies, Albert Einstein College of Medicine, Bronx, NY | Mouse MAb |
| PHF-1 | pSer396/pSer404 | Peter Davies, Albert Einstein College of Medicine, Bronx, NY | Mouse Mab |
| Total Tau Antibodies |  |  |  |
| DA9 | aa102-140 | Peter Davies, Albert Einstein College of Medicine, Bronx, NY | Mouse Mab |
| DA31 | aa150–190 | Peter Davies, Albert Einstein College of Medicine, Bronx, NY | Mouse Mab |
| *Epitope based on human Tau-441 sequence. |  |  |  |

### Cell lines and media

Brain mouse neuroblastoma N2a cells were purchased from American Type Culture Collection (ATCC #CRL-2266, Manassas, VA, USA) and were grown as recommended by the manufacturer. The cells were grown at 1:1 Dulbecco’s modified Eagle medium: reduced serum Eagle’s minimum essential media (DMEM: Opti-MEM) supplemented with 5% FBS (Thermo-Fisher), 100 units/mL penicillin and 0.1 mg/mL streptomycin. Cells were incubated at 37°C in a humidified 5% CO_2_-containing atmosphere.

### Primary Cerebrocortical neuronal cultures

Rat primary cerebrocortical neuronal culture (CTX; Thermofisher; Cat. No. A10840) harvested from a homogenized pool of day ten Sprague–Dawley rat brains and plated on poly-L-lysine-coated (0.01% (w/v)) 12-well culture plates (Erie Scientific, Portsmouth, NH, USA), similar to previously described methods[102] at a density of 4.36 × 10^5^ cells/ml. Cultures were grown in Neurobasal® media (Thermo Fisher), supplemented with 1% B-27 (Thermo Fisher), one mM Glutamine (Thermo Fisher) and incubated at 37°C in a humidified 5% CO_2_-containing atmosphere. The medium was replaced every three days.

### Cell treatments

For N2a cell culture treatments, complete media was replaced with serum-free DMEM media. For CTX primary cultures, all experiments were performed after ten days in culture, and the media was replaced with Neurobasal® media supplemented with 0.5% B-27. For both CTX and N2a culture, SNJ-1945 (S, 100 µM) and Z-DCB (Z, 60 µM) were added to all experimental conditions before the treatment for 1h. This was followed by treatment with okadaic acid (OA; 100 nM) for 24h followed by protein kinase inhibitors for 6h. The protein kinase inhibitors used included: K252a (10 µM), AR-A014418 (60 µM), A-1070722 (60 µM), Saracatinib (100 µM), LiCl (5 mM) TBB (30 µM), EGTA (five mM), Roscovitine (60 µM), STS (0.5 µM), CsA (60 µM) (if added) (Table 1).

### Cell Lysate Collection and Preparation

The culture lysate harvesting for N2a cells and CTX culture were identical. After the treatment, conditioned media were collected from each well and added into separate tubes on ice and centrifuged at 10,000 x g for 10 min at 4°C. Lysis buffer was added to the attached cells on the 12-well plates (100 µl per well). The Triton-X lysis buffer included: 1mM DTT, 1% phosphatase inhibitors (Sigma), 1% Mini-Complete protease inhibitor cocktail tablet (Roche Biochemicals), and 1% Triton X-100. The attached cells were then scraped down into the lysis buffer and collected into separate 1.5 ml Eppendorf tubes. The insoluble pellets from the conditioned culture media were combined with the lysed cells in the lysis buffer. The cell lysates were incubated for 90 minutes at 4°C and then centrifuged at 15,000 rpm for 15 minutes to remove cell debris.

### SDS–PAGE and Western blotting

Protein concentrations of cell lysates were determined by bicinchoninic acid microprotein assays (Pierce Inc., Rockford, IL, USA) against albumin standards. Equal protein samples (20 μg) were prepared for SDS–PAGE in 8x loading buffer containing 0.25 M Tris (pH 6.8), two mM DTT, 8% SDS, and 0.02% bromophenol blue. Each sample was subjected to SDS–PAGE electrophoresis on a 4-20% precast-gels (Bio-Rad) and then transferred on to PVDF membranes. The membranes were blocked in 5% milk for 1h and then incubated with primary antibodies (1/1000) overnight. The secondary antibodies (Amersham Biosciences, UK, 1/10,000) anti-rabbit or anti-mouse IgG conjugated with alkaline phosphatase (Amersham, Piscataway, NJ, USA), were then added for 1h at room temperature. The blots were then washed with TBST, and immunoreactive bands were visualized by developing with biotin, avidin-conjugated alkaline phosphatase, nitro blue tetrazolium, and 5-bromo-4-chloro-3-indolyl phosphate (BCIT/NBT) developer (KPL, Gaithersburg, MD, USA). A 250 kDa to 14 kDa rainbow molecular weight marker (RPN800E, GE Healthcare, Bio-Sciences, Pittsburgh, PA, USA) was loaded in the first well of the electrophoretic gel to estimate the molecular weight of each band. Quantitative evaluation of protein levels was performed via computer-assisted densitometric scanning (NIH ImageJ, version 1.6 software).

### Statistical Analysis

Statistical analysis was performed with one-way ANOVA Tukey’s Test. For multiple comparisons, one-way ANOVA followed by the Bonferroni’s post-hoc test was performed. *p<0.05, **p<0.01, ***p<0.001, **** p<0.0001, ns: non-significant. GraphPad Prism 8.0 (GraphPad, La Jolla, CA).

## Results

Okadaic acid (OA), a potent PP2A/PP1 inhibitor, is known to induce tau hyperphosphorylation and aggregation (43, 44). To establish our tauopathy-relevant cell model, mouse neuroblastoma N2a cells were treated with okadaic acid (OA) (100 nM) to induce tau hyperphosphorylation and oligomerization for 6h and 24h (Figure 1a, b). This specific concentration of OA was selected based on other studies that used similar concentrations optimized on neuronal cell culture (38, 44–46).

**Figure 1.**
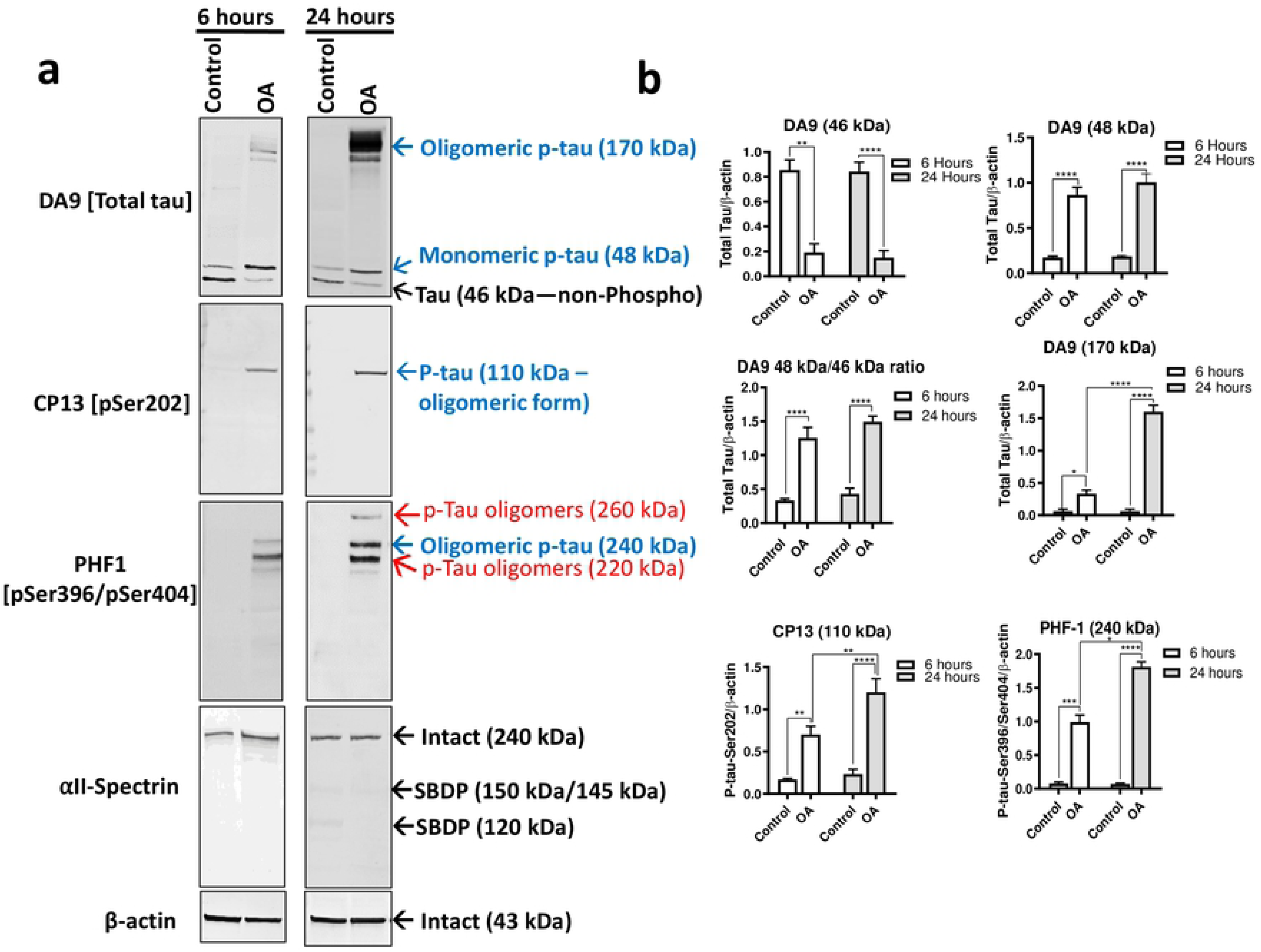
OA induced tau hyperphosphorylation and oligomerization at different time points in mouse neuroblastoma N2a cells. (a). Immunoblots of N2a cells extracted protein (20 µg) using total and phospho-tau antibodies: DA9 (a.a. 102-140), CP13 (pSer202), and PHF-1 (pSer396/pSer404). αII-Spectrin antibody was used to assess neuronal apoptotic pathway activation through monitoring intact spectrin (240 kDa), SBDP150/145 (calpain activation), and SBDP120 (caspase-3 activation). Different tau species are pointed with colored arrows. Blue arrows present monomeric p-tau (48 kDa), and oligomeric p-tau (110 kDa, 170 kDa, and 240 kDa). Red arrows on PHF-1 points on two minor bands of oligomeric p-tau (220 kDa and 260 kDa). Black arrows show non-phospho tau band (46 kDa). SNJ-1945 (abbreviated as S; a calpain inhibitor, 100 μM) and Z-DCB (abbreviated as Z; a caspase-3 inhibitor, 60 μM) were added for all experimental conditions for 1h before the treatment with OA (100 nM) for 6h or 24h, to prevent apoptosis-mediated proteolysis of tau and αII-Spectrin. A reverse time course followed OA treatment, and all cells were collected at the same time and conditions. (b). Immunoblots quantification. All data are normalized to β-actin and are expressed as a percentage of control. Data are presented as ± SEM for n=3. Statistical analysis was performed with one-way ANOVA. For multiple comparisons, one-way ANOVA followed by the Bonferroni’s post-hoc test was performed. *p<0.05, **p<0.01, ***p<0.001, ****p<0.0001 and ns: non-significant. Full-length blots are presented in (**Supplementary Figure 4**).

Since OA is known to induce apoptosis (47), cell-permeable calpain (SNJ-1945) and caspase-3 (Z-DCB) inhibitors were included in all of our experimental conditions in order to eliminate modifications resulting from cell metabolism/health (48, 49). To assess cell viability, caspase-3, and calpain activation, the samples were probed for for αII-spectrin integrity. αII-spectrin is a key substrate for cysteine proteases associated with necrosis (calpain) and apoptotic (caspase-3) cell death (50). Cleavage of αII-spectrin by calpain produces major spectrin break down products (SBDP) of molecular weight 150 kDa (SBDP150) and 145 kDa (SBDP145), while caspase-3 activation produces major cleavage product of 120 kDa (SBDP120) detectable by Western blotting (50, 51). Our control samples probed with αII-spectrin detected only a high molecular weight 240 kDa band (intact αII-spectrin); while SBDPs were absent, suggesting a healthy metabolism and neuronal culture (Fig 1a). Western blots were analyzed with total tau monoclonal antibody DA9 (a.a. 102-140) and monoclonal phospho-tau antibodies including CP13 (pSer202) and PHF-1 (pSer396/pSer404) (Table 1). β-actin was probed to evaluate the evenness of loading the protein extracts. Untreated control showed that the total tau antibody DA9 detected tau protein bands at 46 kDa and 48 kDa at 6h and 24h (Figure 1a). The intensity of the band at 46 kDa was detected at higher levels compared to the band at 48 kDa in control samples. (Figure 1a).

Treatment with OA (100 nM) for 6h and 24h showed a dramatic decrease in levels of the 46 kDa and increased levels of 48 kDa with DA9 antibody. One might presume that the 46 kDa and 48 kDa bands are different tau isoforms. However, how OA treatment affected these bands suggests that they are representative of phosphorylated (p-tau) and non-phosphorylated tau (tau) rather than being tau isoforms. Thus, in our study,the 46 kDa was assigned as tau and the 48 kDa as p-tau.

Additionally, treatment with OA showed high molecular weight (HMW) band clusters residing at 170 kDa probed with DA9 antibody (a.a 102-145) for 6h (p<0.05) and at 24h (p<0.0005). These (HMW) bands may represent the formation of tau oligomers as they were not observed in control cells and only with OA-treated cells. It has been reported in a recent study that treatment with OA in human neuroblastoma SH-SY5Y cells induced tau phosphorylation and oligomerization (44). Another study showed that a local injection of OA in mice induces tau phosphorylation and aggregation in different anatomical brain regions (43). Because the tau phosphorylation and formation of HMW bands were observed relatively at higher levels with OA treatment for 24h compared to the 6h, the 24h treatment was selected as our tauopathy model (Figure 1a, 1b).

On the other hand, in our cell culture experimental conditions, treatment with OA (100 nM) for less than 6h did not show any detectable tau bands with CP13 and PHF-1 (data not shown). It is well-known that OA induces apoptosis in human neuroblastoma cells, mouse neuroblastoma, and rat cerebellum neurons (47). Thus, the time points were not increased beyond 24h of treatment to avoid tau phosphorylation modifications resulting from proteolysis and neural death.

Probing with CP13 (pSer202) antibody did not show any detectable bands of tau protein in control samples (Figure 1a). This result indicates that endogenous phosphorylation of tau at Ser202 site is low under normal growth conditions. However, with OA treatment, CP13 showed HMW band formed at 110 kDa (x2 size of monomeric tau) with 6h and 24h (Figure 1a, 1b). Probing with PHF-1 antibody (pSer396/pSer404) did not show any tau band with control samples (Figure 1a, 1b). Treatment with OA for 6h and 24h showed HMW cluster of bands at 220 kDa, 240 kDa, and 260 kDa with PHF-1 (Figure 1a, 1b). Notably, the 260 kDa band (red arrow) (Figure 1a) was only detectable with OA treatment for 24h (PHF-1).

Low molecular weight monomeric tau (LMW-MT) bands were not detected with either CP13 or PHF-1. It should be noted that the DA9 antibody recognizes total tau epitopes from aa. 102-140. Thus, to identify the same 48 kDa tau species detected with DA9, the phospho-tau antibody needs to recognize the same epitope. It was assumed that LMW-MT might be either phosphorylated at sites other than Ser202/Ser396/Ser404, and LMW-MT oligomerized into the different HMW tau species detected at 110 kDa, 170 kDa, 220 kDa, 240 kDa, and 260 kDa. Indeed, using RZ3(pThr231) and AT270(pThr181), LMW-MT at 48 kDa and 55 kDa was detected with OA treatment; respectively **(Supplementary Figure 1a, left panel, OA lane)**.

Taken together, these data strongly suggest that OA treatment caused protein phosphatase inhibition inducing the formation of LMW and HMW tau bands, immunoreactive at pSer202 (CP13, 110 kDa), pSer396/pSer404 (PHF-1, 220/240/260 kDa), RZ3 (pThr231, 48 kDa) and AT270 (pThr181, 55 kDa). Based on the molecular weight of each species, the immunoreactivity with tau antibodies solidifies the notion of tau hyperphosphorylation and oligomerization.

### Screening of tau kinase inhibitors on OA-induced tau hyperphosphorylation and oligomerization in N2a cells

To screen for protein kinase inhibitors as drug candidates for inhibition of OA-induced hyperphosphorylation and oligomerization, mouse neuroblastoma N2a cells were pre-treated with OA for 24h followed by treatment with protein kinase inhibitors for 6h. The positive control included only OA treated cells for 24h. Protein kinase inhibitors used included: LiCl (10 mM), AR-A014418 (AR) (60 µM), A-1070722 (A107)(60 µM), K252a (10 µM), STS (0.5 µM) 4,5,6,7-tetrabromobenzotriazole (TBB) (60 µM), Roscovitine (60 µM), Saracatinib (100 µM), cyclosporine A (CsA) (60 µM), and EGTA (5 mM) (Table 1; Figure 2a, 2b). All conditions were pre-treated with SNJ-1945 (calpain inhibitor, abbreviated as S; 60 µM) and Z-DCB (caspase inhibitor, abbreviated as Z; 100 µM) to minimize apoptotic pathways activation (calpain and caspase-mediated proteolysis) (48, 52) (Table 1). To assess cell integrity, the αII-spectrin antibody was used to monitor intact-240 kDa, SBDP150, and SBDP120 that are representatives of apoptosis, necrosis, calpain, and caspase activation, respectively.

**Figure 2.**
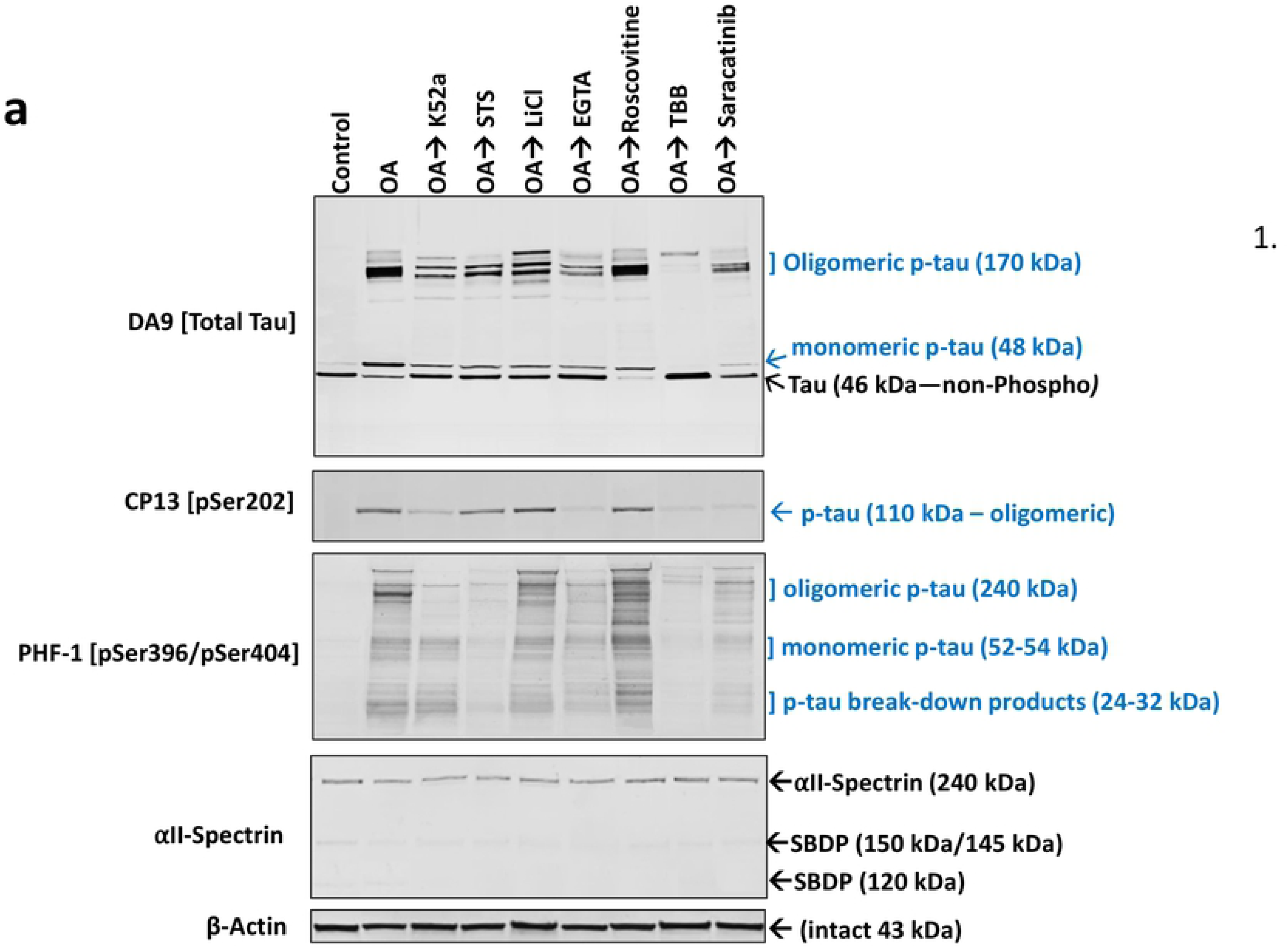
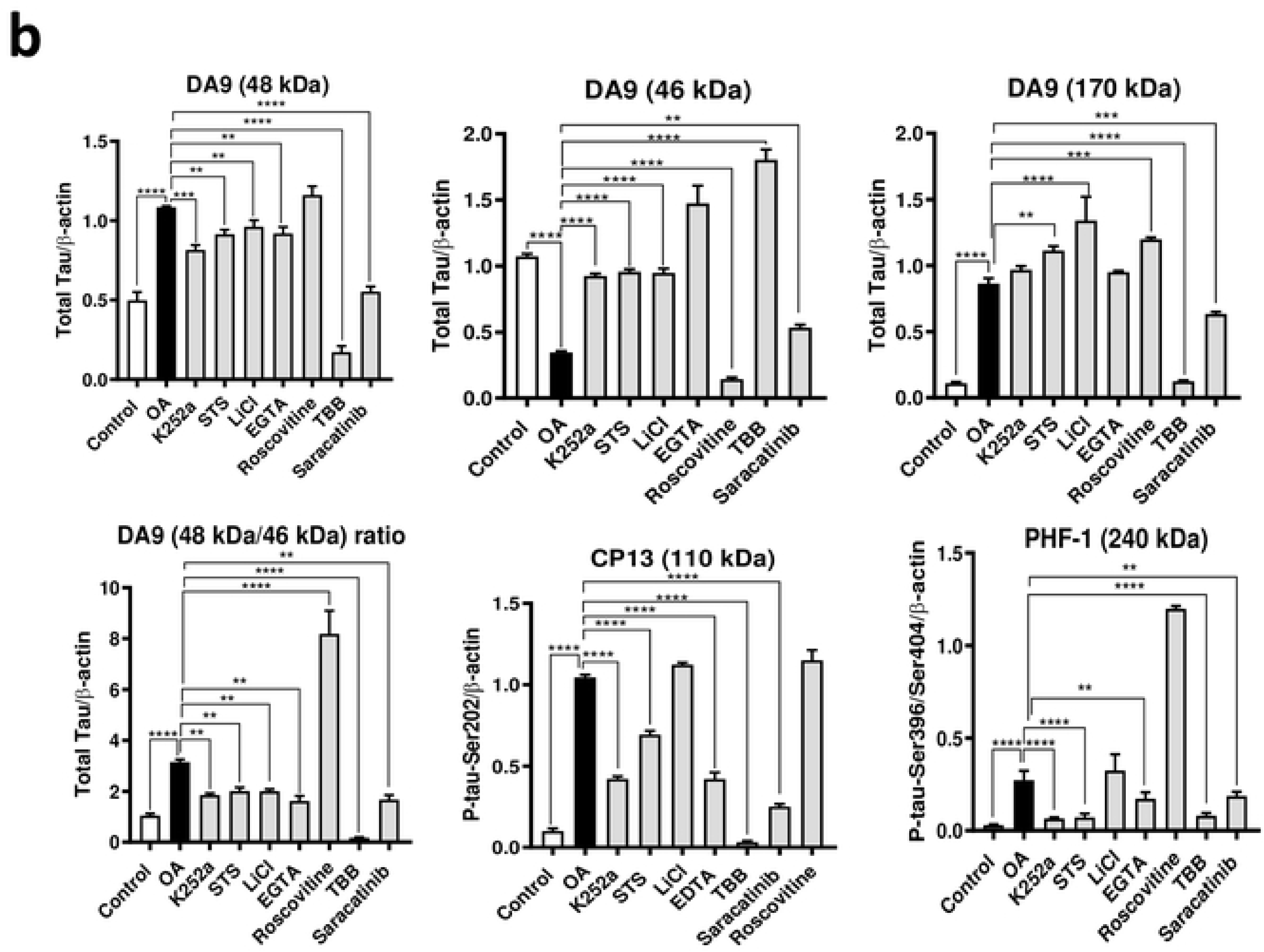
Screening of protein kinase inhibitors on OA-induced Tau hyperphosphorylation and oligomerization in N2a cells. (a). Immunoblots of N2a cells extracted protein (20 µg) using phospho-tau antibodies (CP13, PHF-1), total tau (DA9), and αII-Spectrin. αII-Spectrin was probed to assess neuronal cell injury monitored with SBDP145/150 and SBDP120. Kinase inhibitors effect on OA-induced tau bands (100 nM) was monitored by evaluating the levels of monomeric (48 kDa) and oligomeric p-tau immunoreactivity (110 kDa, 170 kDa, and 240 kDa; blue arrows), total tau, and non-phospho tau (46 kDa; black arrows). Phosphorylated tau break-down products are shown with PHF-1 immunoblot. For all experimental conditions, S (a calpain inhibitor) and Z (a caspase-3 inhibitor) were added for 1h to before the addition of OA for 24h followed by 6h incubation with the kinase inhibitors. The concentrations used for each protein kinase inhibitor are mentioned in materials and methods, cell treatment section. β-actin was probed as a loading control. All experimental conditions were collected and analyzed at the same time. (b). Immunoblots quantification. All data are normalized to β-actin and are expressed as a percentage of control. Data are presented as ± SEM for n=3. Statistical analysis was performed with one-way ANOVA. For multiple comparisons, one-way ANOVA followed by the Bonferroni’s post-hoc test was performed. *p<0.05, **p<0.01, ***p<0.001, ****p<0.0001 and ns: non-significant. Full-length blots are presented in (**Supplementary Figure 5**).

### Casein kinase II (CKII) inhibitor: 4,5,6,7-tetrabromobenzotriazole (TBB)

Since aberrant CKII has been reported in AD (53), TBB, a cell-permeable CKII inhibitor, was selected for the study. Total tau DA9 showed that TBB abolished the 48 kDa band (p-tau) and the HMW 170 kDa band (tau oligomers), and significantly increased (p<0.0001) levels of the 46 kDa (non-phospho tau) by 85%, compared to OA treatment alone (Figure 2a, 2b, Table 3). CP13 (pSer202) antibody showed that TBB eliminated the OA-induced 110 kDa band (oligomeric p-tau). Similarly, PHF-1 antibody (pSer396/pSer404) showed that TBB fully inhibited the formation of 240 kDa (HMW bands-oligomeric tau) (Figure 2a, 2b, Table 3). As a selective casein kinase II (CKII) inhibitor, TBB showed robustness in inhibiting both OA-induced tau hyperphosphorylation and oligomerization. Thus, the aim was to evaluate the TBB dose-response effect on OA-induced tau hyperphosphorylation and oligomerization in N2a neuronal culture. To achieve this aim, N2a cells were treated with OA for 24h followed by treatment with various concentrations of TBB (10 nM, 30 nM, 100 nM, 300 nM, 1 µM, 3 µM, 10 µM, and 30 µM) for 6h (Figure 3a, 3b). The result shows that treatment with ten micromolars of TBB resulted in 50% reduction of the 110 kDa (oligomeric p-tau form; CP13), 48 kDa, and 170 kDa bands (monomeric and oligomeric p-tau, DA9) (Figure 3a, 3b). Increasing the concentration of TBB up to 30 µM caused 90% reduction of 48 kDa (monomeric p-tau, DA9), 170 kDa (oligomeric p-tau, DA9) and 110 kDa (oligomeric tau form, CP13) (Figure 3a, 3b). As for assessing neuronal culture integrity, the intact αII-spectrin band was detected at 240 kDa, and no SBDP150/145 or SBDP120 was observed with the TBB treated conditions suggesting a healthy culture.

**Figure 3.**
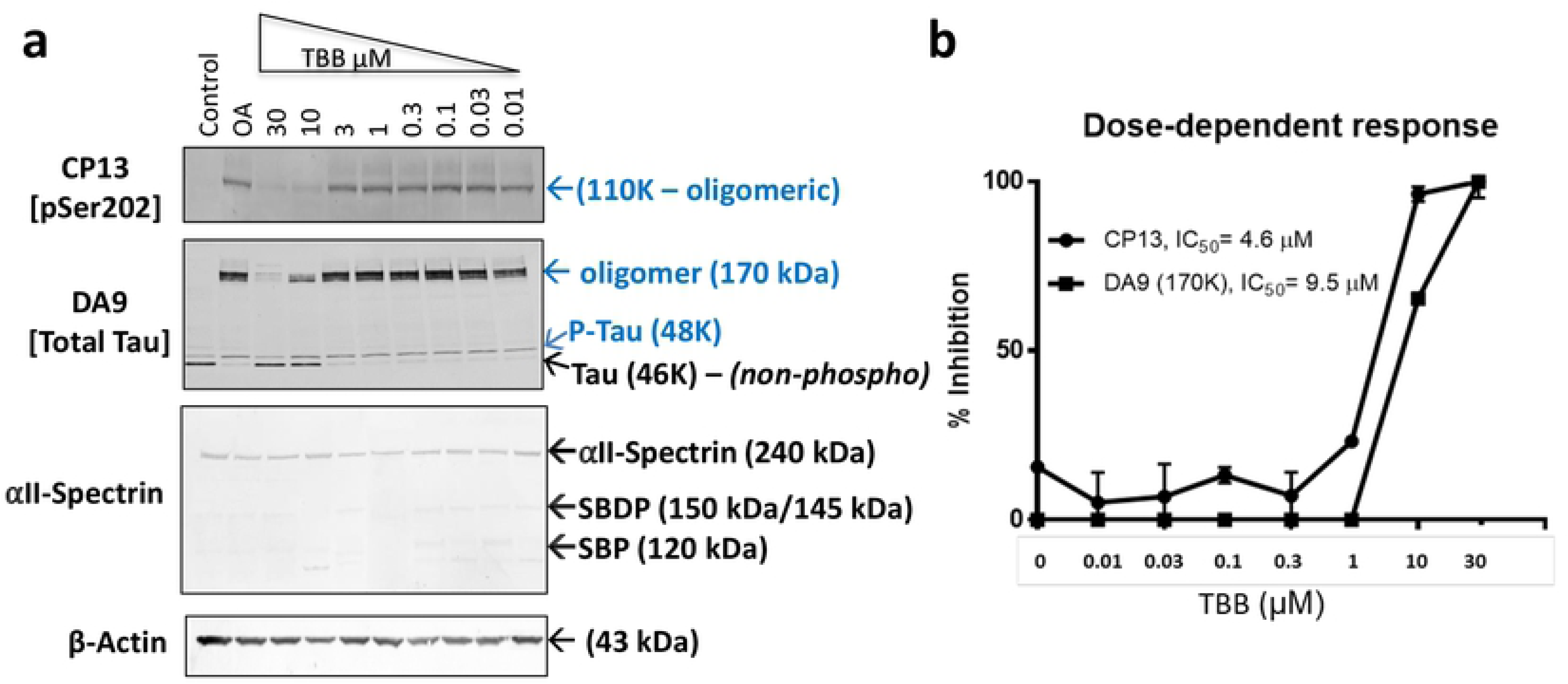
Dose-response of TBB on OA-induced tau hyperphosphorylation and oligomerization in N2a cells. N2a cells were pre-treated with OA for 24h followed by treatment with different concentrations of TBB for 6h, as indicated in the figure. (a) Immunoblots of cell extracted proteins using phospho-tau antibodies, including CP13 (pSer202), and total tau DA9 (a.a. 102-140). Blue arrows represent monomeric and oligomeric p-tau (48 kDa, 110 kDa, and 170 kDa). αII-Spectrin antibody used to monitor SBDPs with the increasing concentrations of TBB. The β-actin antibody was used as a loading control. All conditions included SNJ-1945 (calpain inhibitor) and Z-DCB (caspase inhibitor). (b) TBB dose-response treatment line chart. TBB concentration (in micromolar) is shown on the X-axis, and the inhibition percentage is presented on the Y-axis. The control sample values were designated as the standard response. The X-axis concentration values are logarithm-transformed to fit a straight line. The half maximal inhibitory concentration (IC50) was used to measure the effectiveness of TBB in inhibiting OA-induced tau hyperphosphorylation and oligomerization. GraphPad Prism was used to calculate the IC50 (for DA9 and CP13 antibodies) and are presented in the figure. The statistical analysis was performed with one-way ANOVA, followed by Bonferroni’s post-hoc test. *p<0.05, **p<0.01, ***p<0.001. Data are presented as ± SEM for n=3. Full-length blots are presented in (**Supplementary Figure 6**).

**Table 3.** Composite effects of kinase inhibitors on OA-induced Tau hyperphosphorylation in N2a cells.

|  |  |  |  |
| --- | --- | --- | --- |
| Inhibitor | Inhibition % |  |  |
|  | CP13<br>(110 kDa) | PHF-1<br>(240 kDa) | DA9<br>(170 kDa) |
| K252a | 62 | 70 | 40 |
| STS | 32 | 82 | -10 |
| LiCl | -14 | -9 | -12 |
| EGTA | 90 | 85 | 55 |
| CsA | 98 | 90 | **100** |
| Roscovitine | -11 | -53 | -22 |
| **TBB** | **100** | **100** | **100** |
| Saracatinib | 70.5 | 46 | 45 |
| A107 | 23 | 0 | 13 |
| AR | 0 | 0 | 36 |

A negative sign correspond to an adverse effect. Bold corresponds to maximal inhibition at the tested epitope.

### Calcineurin Inhibitor: Cyclosporin A (CsA)

Cyclosporin A (CsA) has been reported to inhibit calcineurin phosphatase activity (PP3) and CaMKII by blocking the Ca++ mitochondrial permeability (54) (Table 1). Thus, CsA was selected in this study as a calcium-dependent kinase inhibitor to assess its effect on OA-induced tau hyperphosphorylation and oligomerization. Notably, CsA abolished the 48 kDa and 170 kDa monomeric and oligomeric p-tau of DA9, respectively (**Supplementary Figure 1a**). Moreover, CsA significantly reduced the protein band at 110 kDa of CP13 (oligomeric p-tau band; p<0.0001), and 240 kDa of PHF-1 (oligomeric p-tau band) (**Supplementary Figure 1a, b**, Table 3). Phosphorylation sites at human tau threonine 181 and 231 have been shown to differentiate AD patient from a control subject (35, 55). Hence, RZ3 (pT231) and AT270 (pT181) antibodies were used to study these sites.

RZ3 antibody showed a complete reduction of the 48 kDa band (monomeric p-tau) when cells were treated with CsA (**Supplementary Figure 1a, b**). As for AT270 antibody, OA-treated samples showed a band detected at 55 kDa (monomeric p-tau) which was abolished when neurons were treated with CsA. The 48 kDa and 55 kDa OA-induced bands detected by RZ3 and AT270, respectively, indicate that tau can have various levels of oligomerization, or can still be monomeric, depending upon the phosphorylation site tested. β-actin protein levels remained even in all experimental conditions. As for testing neuronal injury and apoptotic pathway activation, αII-spectrin blotting did not detect any significant changes of the 240 kDa band (intact form). Moreover, SBDP150/145 or SBDP120 immunoreactive bands were not detected in all of the treated samples, indicative of a healthy metabolism.

### Calcium chelator: EGTA

Another calcium-dependent kinase inhibitor, EGTA, was used as a calcium-chelating agent. EGTA has a lower binding affinity for Mg^++^ relative to EDTA, making it more selective for Ca^++^ ions (56). Total tau DA9 showed that EGTA treatment (with calpain and caspase inhibitors; S+Z) caused 25% reduction of the 48 kDa band (p-tau), 85% increase of the 46 kDa band (non-phospho-tau) and 55% reduction of the 170 kDa band (oligomeric p-tau form), compared to OA treatment alone (Figure 2a, 2b and Table 3). Additionally, EGTA caused 90% reduction of 110 kDa band (oligomeric tau; CP13) and 85% reduction of 240 kDa (oligomeric tau; PHF-1) (Figure 2a, 2b and Table 3). As for apoptotic pathway activation, αII-spectrin antibody did not show any effect on the 240 kDa band (intact form), and the SBDP150/145 or SBDP120 bands were not detected with EGTA treatment.

### Glycogen synthase kinase-3 (GSK-3) inhibitors: LiCl, A-1070722, and AR-1014418

Tau is a substrate of Glycogen synthase kinase-3 (GSK-3)(57), and p-tau phosphorylation and oligomerization could be inhibited by GSK-3 inhibition. To test this hypothesis, the effects of small molecule GSK-3 inhibitors, LiCl, A-1070722 (abbreviated as A-107), and AR-1014418 (abbreviated as AR) were examined on OA-induced tau hyperphosphorylation and oligomerization. Surprisingly, LiCl showed an opposite effect in N2a cell treatment by increasing levels of 110 kDa band (oligomeric form, CP13; −14%) and levels of 240 kDa band (oligomeric p-tau, PHF-1; −9%) (Figure 2a, 2b and Table 3). As for total tau DA9 antibody, LiCl also showed an opposite effect by increasing the 48/46 kDa (p-tau/non-phospho-tau) band ratio by 20%, and the 170 kDa oligomeric tau band by 12%.

AR, a thiazole class inhibitor, was shown to decrease insoluble p-tau in the brain stem of transgenic mice overexpressing a mutant human tau protein (58). In our experimental design, AR did not show a statistically significant effect on the 240 kDa band (oligomeric p-tau; PHF-1) or 110 kDa band (oligomeric p-tau form; CP13) (**Supplementary Figure 2a, b**, Table 3). Moreover, probing with total tau DA9 showed that AR treatment caused an adverse effect by increasing the 48 kDa/46 kDa ratio (monomeric tau form; −50%) and reducing the 170 kDa oligomeric form band by 36% (**Supplementary Figure 2a, b**, Table 3).

Another potent GSK-3 inhibitor, A-107 (Ki=0.6 nM for GSKα and GSK-3β) (59), was selected for the study. OA followed by A-107 treatment showed 23% reduction of 110 kDa oligomeric form band (CP13) and non-significant but partial reduction of the 240 kDa oligomeric p-tau form (PHF-1) compared to OA treatment alone (**Supplementary Figure 2a, b**, Table 3). Probing with total tau DA9 showed with A-107 treatment, a 13% reduction of 170 kDa band (DA9) and did not show a statistically significant effect on the 48 kDa band (**Supplementary Figure 2a, b**, Table 3). As for caspase-3, calpain, and cell injury activation, αII-spectrin did not show SBDP 150/145 or SBDP120 post-treatment, indicative of a healthy neuronal culture.

### Src/Fyn kinase inhibitor: Saracatinib

Saracatinib is an inhibitor of the Src/abl kinase family, developed initially for several types of cancer but withdrawn for the lack of effectiveness (60). However, Saracatinib is also a potent inhibitor of Fyn kinase, which is linked to tau (16). Fyn has been reported to phosphorylate dendritic tau, which allows Fyn to localize to the post-synaptic density (16). In the current study, Saracatinib was selected to investigate the role of Fyn kinase function on the tauopathy-relevant cell-based model. Probing with DA9 (a.a. 102-140) antibody, Saracatinib resulted in 40% reduction in the 48 kDa (monomeric p-tau), 20% increase in 46 kDa (non-phospho tau) bands and 45% reduction of the 170 kDa band (oligomeric p-tau form) (Figure 2a, 2b, Table 3). Saracatinib treatment caused significant reduction (p<0.0001) of the 110 kDa oligomeric p-tau band of CP13 (75%; CP13) and produced 46% reduction in immunoreactivity of the 240 kDa (oligomeric p-tau form band of PHF-1) (Figure 2a, 2b, Table 3). As for assessing cell integrity, intact spectrin (240 kDa) levels remained constant, and SBDP150/145 and SBDP120 levels were not significantly altered, with Saracatinib treatment, compared to control values.

### Pan kinase inhibitor: K252a and STS

K252a is a non-selective cell-permeable protein kinase inhibitor, inhibiting protein kinase C (PKC; IC50=32.9 nM), Ca2+/calmodulin-stimulated phosphodiesterases (IC50=1.3-2.9 μM), serine/threonine protein kinases (IC50=10-20 nM), myosin light-chain kinase (MLCK; Ki=20 nM), receptor tyrosine kinases, and inhibiting the carcinogenic properties of MET oncogene (61, 62). K252a is an analog of staurosporine (STS) and has a broad spectrum of protein kinases inhibition, neuroprotection properties, and improvement in psoriasis in vivo (Table 1) (63). In this study, K252a and STS treatment similarly showed 30% increase in 46 kDa (monomeric non-phosphorylated tau) and 35% decrease at 48 kDa (monomeric p-tau) compared to OA, with DA9 antibody (Figure 2a, 2b, and Table 3). K252a treatment caused 40% reduction of 170 kDa (DA9; oligomeric p-tau) compared to OA treatment alone (average of n=3) (Figure 2a, 2b, and Table 3). For p-tau detection, probing with CP13 antibody showed 60% and 32% reduction in 110 kDa (oligomeric p-tau form) with K252a and STS treatment, correspondingly. PHF-1 showed 70% and 80% reduction in levels of 240 kDa (oligomeric form) with K252a and STS treatment, respectively (Figure 2a, 2b, Table 3). αII-spectrin immunoreactive bands (intact-240 kDa, SBDP150/145, and SBDP120) did not show a statistically significant difference compared to control values. Although STS is known to induce apoptosis, the absence of SBDPs is due to the effect of caspase and calpain inhibitors (S+Z), which in our previous studies have shown to decrease SBDP150/145/120 resulting from STS treatment (64, 65).

### CDK5 inhibitor: Roscovitine

Roscovitine is a cyclin-dependent kinase 5 (CDK5) that acts through direct competition at the ATP-binding site (66). It has been previously shown that tau protein can be a hyperphosphorylated by CDK5 in specific pathological conditions (67–69). To study the role of CDK5 in our tauopathy cell-based-model, Roscovitine was selected to assess its effect on OA-induced tau hyperphosphorylation and oligomerization. Surprisingly, Roscovitine showed an adverse effect by increasing levels of oligomeric tau detected at 170 kDa (−51%; DA9), 110 kDa (−11%; CP13), and 240 kDa (−53%; PHF-1) compared to OA treatment alone (Figure 2a, 2b, Table 3). Roscovitine showed partial but a statistically non-significant decrease in the 48 kDa (monomeric p-tau) band of DA9, compared to OA treatment alone. Moreover, αII-spectrin antibody did not show a statistically significant difference of intact form (240 kDa), SBDP150/145, and SBDP120 compared to control values. β-actin protein levels remained even in all experimental conditions.

### Baseline and OA-induced tau hyperphosphorylation and oligomerization: effects of various kinase inhibitors treatments in rat primary cerebrocortical neuronal (CTX) culture

To further expand our experimental paradigm in a cell-based model suitable for drug candidate screening, the effectiveness of the protein kinase inhibitors was investigated on rat primary cerebrocortical neuronal (CTX) cultures. Our CTX primary culture is fully differentiated neurons, which can provide a model for physiologically relevant cellular events that make neurons uniquely susceptible to disease-associated proteins. Additionally, the use of high-throughput primary culture allowed us to screen multiple drug candidates in a short period, compared to conventional methods and permit the exposure of novel biological concepts to identify new drug targets for therapeutics. Therefore, CTX cells were pre-treated with or without OA 24h (100 nM) (Table 1) followed by treatment with protein kinase inhibitors for 6h. Calpain and caspase-3 inhibitors, SNJ-1945 and Z-DCB respectively, were added to all experimental conditions to prevent cell death-mediated proteolysis of tau as a potential confound. To monitor neuronal culture health and metabolism, the samples were probed with αII-spectrin antibody, and intact form (240 kDa), SBDP150/145, and SBDP120 were quantified and compared to control.

CTX control cultures showed normal cell bodies and healthy neurites, including axons and dendrites. Notably, untreated control samples showed basal levels of phosphorylated tau (67 kDa) detected by total and p-tau antibodies, including: DA31 (a.a.150-190), CP13 (pSer202), RZ3 (pThr231), PHF-1 (pSer396/pSer404), AT8 (pSer202/pThr205), and AT270 (pThr205) (Figure 4a, 4b, lane 1). In agreement with previous reports using immunocytochemistry and western blotting, rat cortical neurons in primary culture showed that tau is physiologically highly phosphorylated (70). Thus, in our experimental design, various kinase inhibitors were tested on basal and OA-induced p-tau to measure their effects in reducing physiological and pathological phosphorylation levels at different epitopes.

**Figure 4.**
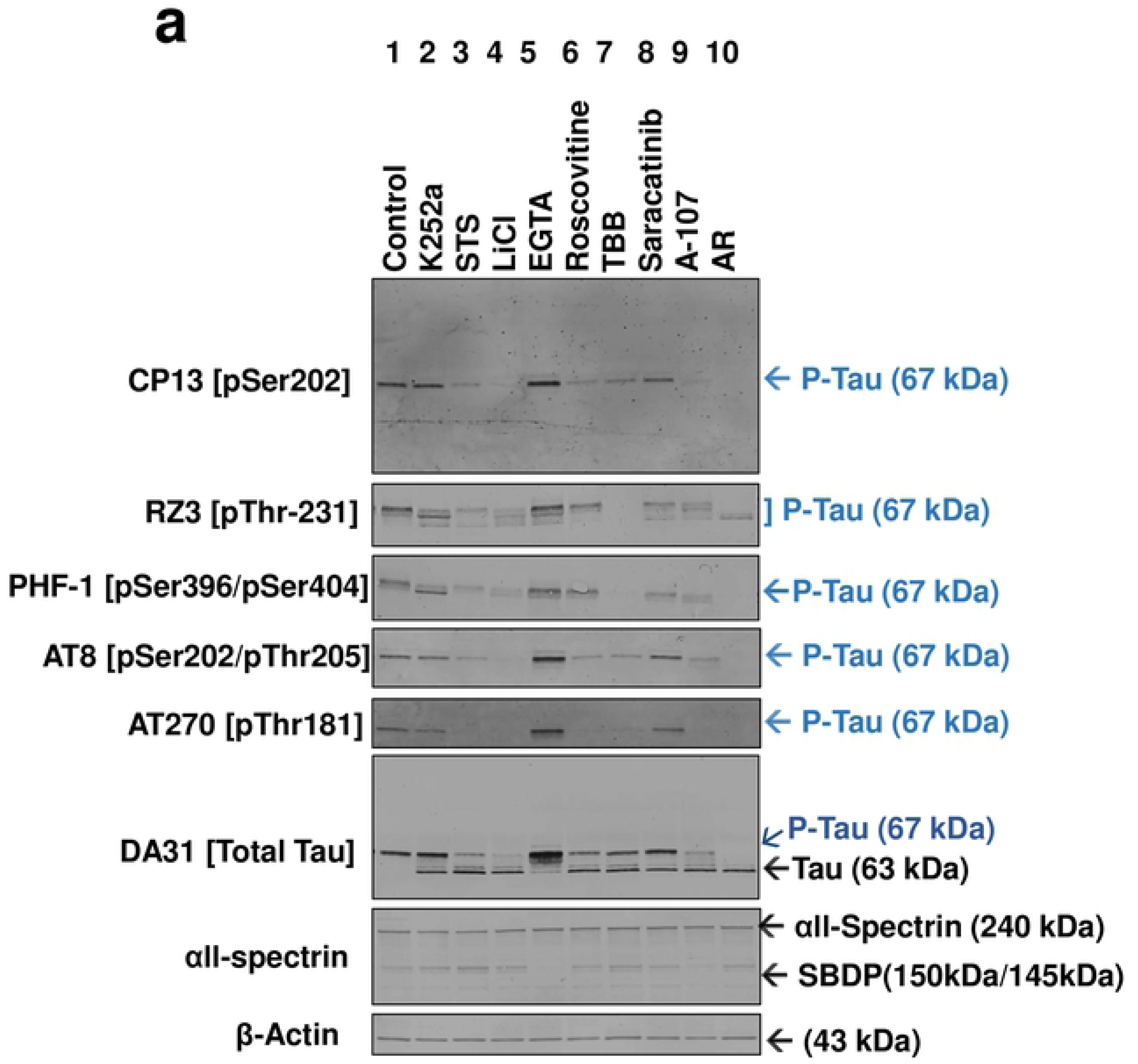
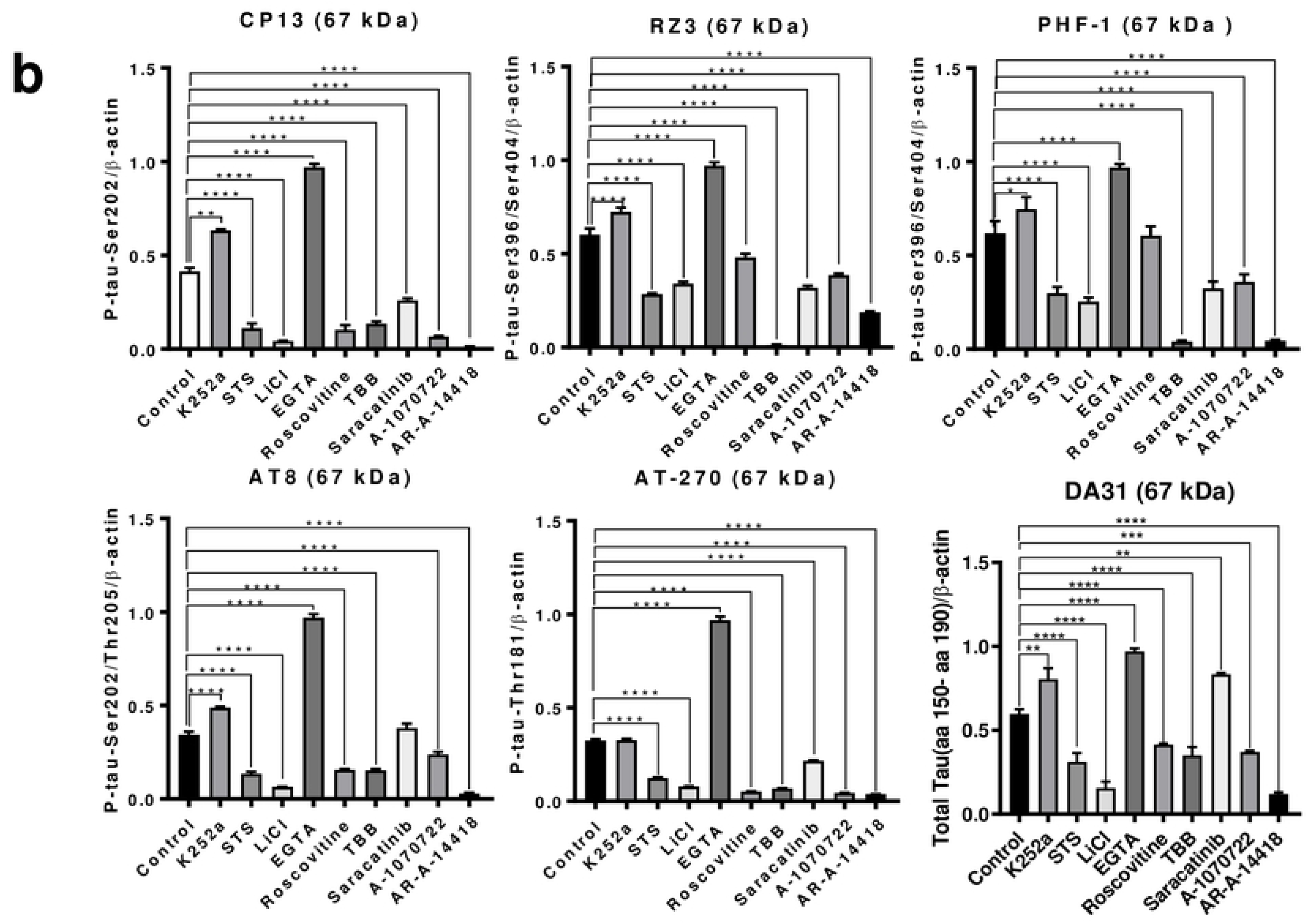
Screening of protein kinase inhibitors on physiologically phosphorylated tau in rat primary cerebrocortical neuronal culture. Rat primary cerebrocortical neuronal differentiated cultures (CTX) at 15 DIVs, were treated various protein kinases inhibitors, including K252a (30 μM), STS (20 μM), LiCl (10 μM), EGTA (5 mM), Roscovitine (60 μM), Saracatinib (100 μM), TBB (30 μM) and A-107 (20 μM), AR (60 μM) for 6h. Calpain and caspase inhibitors (S+Z) were added to all experimental conditions for 1h before the protein kinase inhibitor treatments. Cell lysates were analyzed on western blots using twenty micrograms of protein. (a) Immunoblots of cell lysates analyzed for phosphorylated tau at the epitopes CP13 (pSer202), PHF-1 (pSer396/404), AT8 (pSer202/pThr205), RZ3 (pThr231), and AT270 (pThr181). Total tau was probed with DA31 (a.a. 150-190) antibody. DA31 blot showed two distinctive tau bands (63 kDa, non-phospho tau and 67 kDa, p-tau) following kinase inhibitors treatment. SBDP145/150 and SBDP120 were analyzed with the αII-spectrin antibody. Different lanes are numbered at the top of each label in the figure. (b) Immunoblot quantification of basal tau phosphorylation. Ratios of phospho-epitope levels over β-actin ± SD are represented as a percentage. Statistical analysis was performed with one-way ANOVA. For multiple comparisons, one-way ANOVA followed by the Bonferroni’s post-hoc test was performed. *p<0.05, **p<0.01, ***p<0.001 and ****p<0.0001. n=3 per condition. Full-length blots are presented in (**Supplementary Figure 7**).

Treatment with OA for 24h caused a dramatic increase of 67 kDa band (monomeric p-tau) at multiple phospho-tau epitopes (CP13: 9x, RZ3: 9.8x, PHF-1: 13x, AT8: 3x, and AT270: 10x) (Figure 5a, 5b, lane 2). In contrast to N2a cells, oligomeric forms of tau protein were not observed when tested with total tau (DA31 and DA9), and phospho-tau antibodies in CTX culture. Since the samples were prepared under SDS-reducing conditions, it might be possible that tau oligomers in CTX culture are disrupted, although tau cross-linking by disulfide bonds is not an essential requirement for tau oligomerization (71).

**Figure 5.**
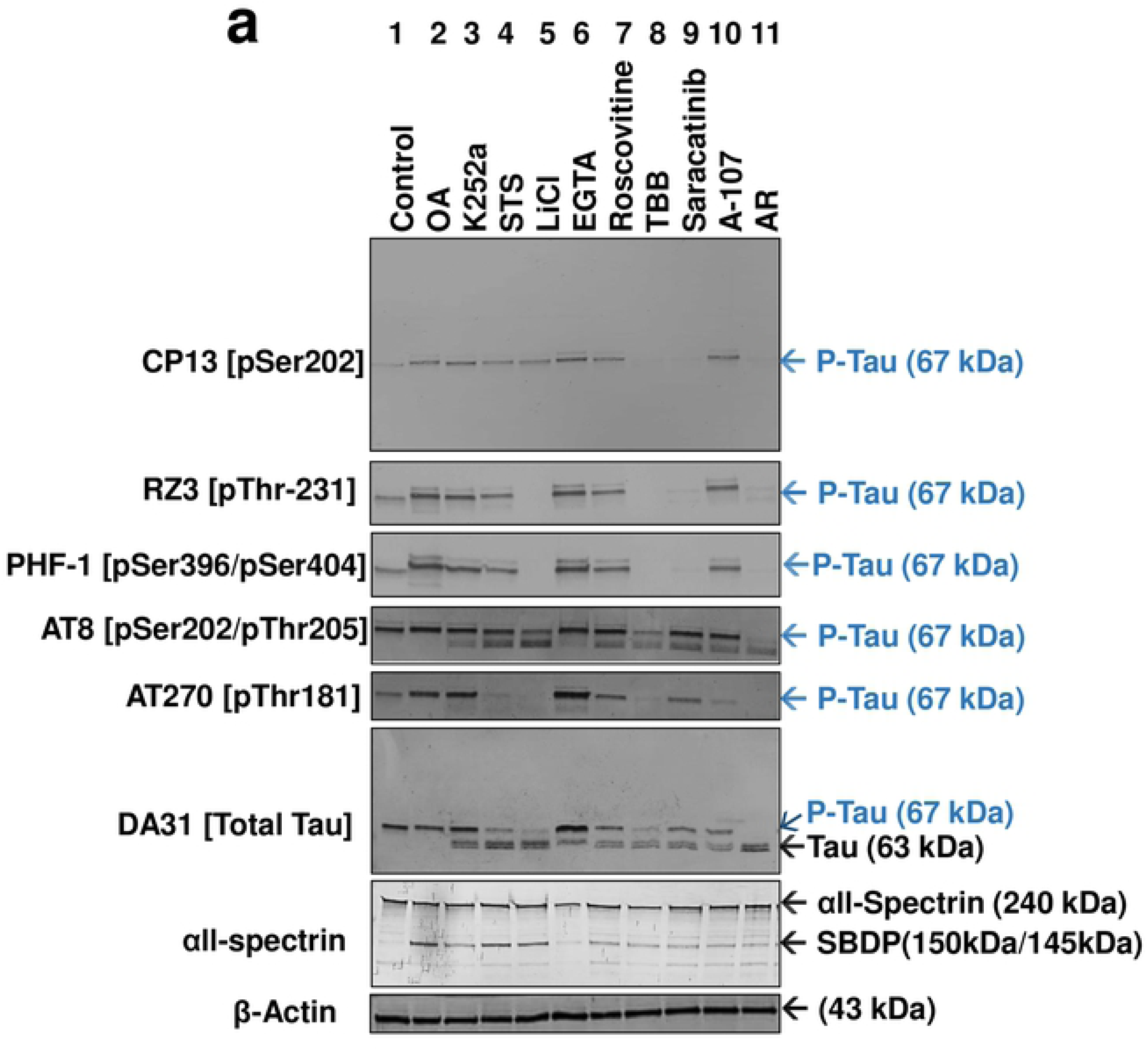
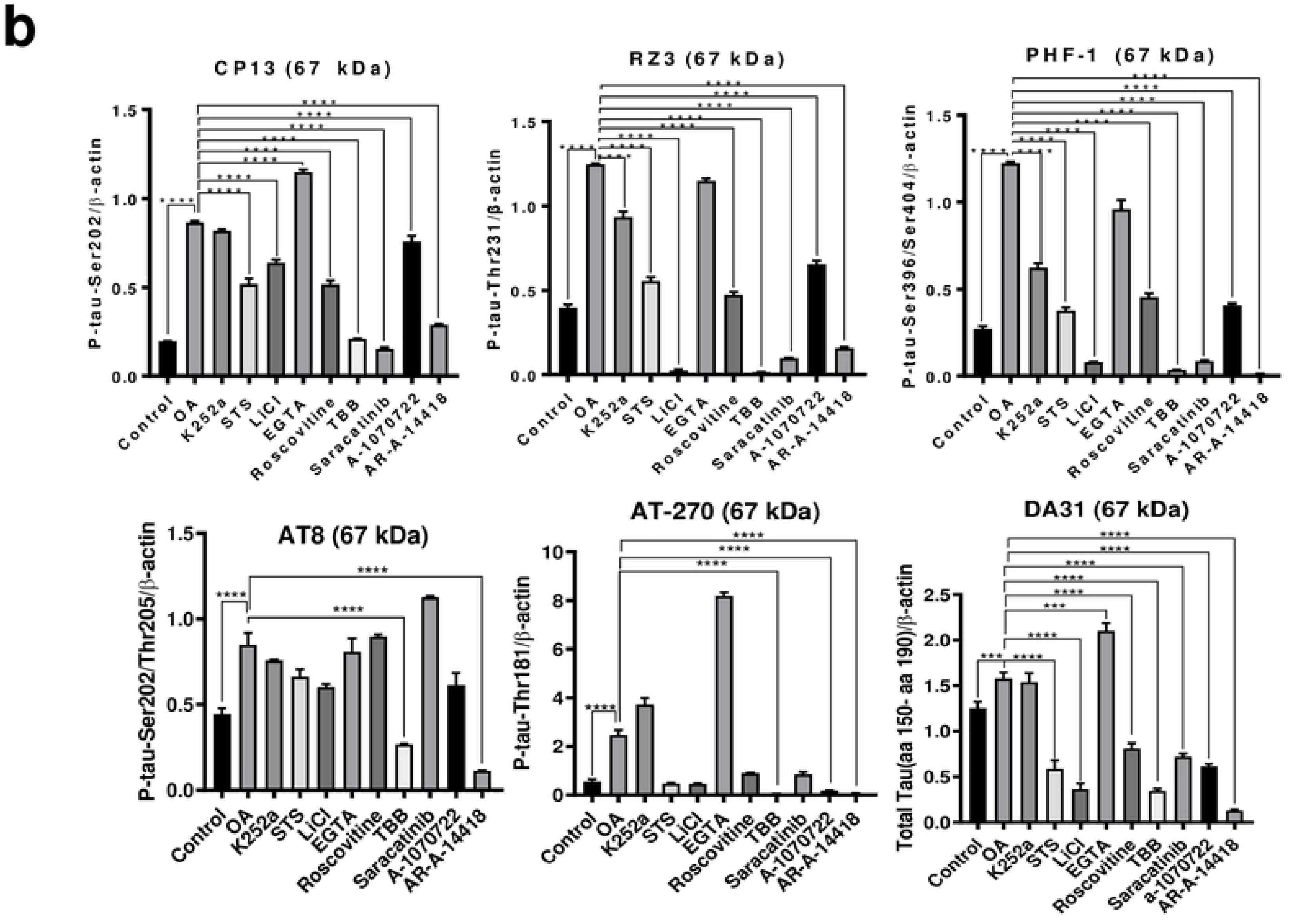
Effect of protein kinase inhibitors on OA-induced tau hyperphosphorylation in rat primary cerebrocortical neuronal culture. Rat primary cerebrocortical neuronal differentiated cultures (CTX) at 15 DIV were treated with OA (100 nM) for 24h followed by protein kinases inhibitors for 6h. The concentrations of kinase inhibitors are the same as the ones mentioned in Figure 4. CTX cultures were treated with S and Z for 1h before any treatment to prevent apoptotic pathway-mediated tau proteolysis. (a). Immunoblots of cell lysates analyzed for phosphorylated tau at the epitopes CP13, PHF-1, AT8, RZ3, AT270. Total tau was probed with DA31 antibody. With DA31 blot, the 63 kDa band is referred to as monomeric non-phospho tau and the 67 kDa as monomeric p-tau species. Spectrin Break down products (SBDPs) were monitored with the αII-spectrin antibody. (b) Immunoblot quantification of OA-induced tau phosphorylation. Ratios of phospho-epitope levels over β-actin ± SD are represented as a percentage. Statistical analysis was performed with one-way ANOVA. For multiple comparisons, one-way ANOVA followed by the Bonferroni’s post hoc test was performed. *p<0.05, **p<0.01, ***p<0.001 and ****p<0.0001. n=3 per condition. Full-length blots are presented in (**Supplementary Figure 8**).

Since the concentration of 30 µM TBB resulted in at least 90% inhibition in N2a cells, the same concentration was used for CTX culture. Treating CTX culture with TBB reduced basal and OA-induced tau phosphorylation (67 kDa) at CP13 (-OA: 91%, +OA: 98%), RZ3 (-OA: 100%, +OA: 100%), PHF-1 (-OA: 100%, +OA: 100%), AT8 (-OA: 91%, +OA: 100%), and AT270 (-OA: 100%, +OA: 100%) compared to OA treatment alone (Figure 4a, 4b, lane 7 and Figure 5a, 5b, lane 8, Table 4). Total tau DA31 (a.a. 150-190) antibody detected immunoreactive bands at 63 kDa and 67 kDa with the different kinase inhibitor treatments. The decreased electrophoretic mobility of the 63 kDa might correspond to the lower levels of p-tau protein induced by the protein kinase inhibitors; thus, this band was assigned as non-phospho-tau. TBB treatment caused a reduction of the phospho-tau band at 67 kDa (-OA: 41%, +OA: 91%), and an increase of non-phospho tau band at 63 kDa (-OA: +53%, +OA: +81%) (Figure 4a, 4b, lane 7 and Figure 5a, 5b, lane 8, Table 4).

**Table 4.** Composite effects of kinase inhibitors on basal and OA-induced Tau hyperphosphorylation in rat primary cerebrocortical neuronal cells.

A negative sign correspond to an adverse effect. Bold corresponds to maximal inhibition at the tested epitope.
|  |  |  |  |  |  |  |  |
| --- | --- | --- | --- | --- | --- | --- | --- |
| Inhibitor |  | Inhibition % |  |  |  |  |  |
|  |  | CP13 | RZ3 | PHF-1 | AT8 | AT270 | DA31 |
| K252a | - OA | 35 | 45 | 61 | 10 | 17 | 41 |
|  | + OA | 41 | 37 | 63 | 0 | 0 | 45 |
| STS | - OA | 86 | 83 | 55 | 88 | **100** | 79 |
|  | + OA | 63 | 81 | 89 | 12 | **100** | 77 |
| LiCl | - OA | 94 | 89 | 98 | **100** | **100** | 93 |
|  | + OA | 50 | 100 | 100 | 81 | **100** | 96 |
| EGTA | - OA | -51 | -63 | -22 | -64 | -68 | -73 |
|  | + OA | -12 | -22 | -13 | -5 | -63 | -69 |
| Roscovitine | - OA | 85 | 42 | 63 | 91 | **100** | 32 |
|  | + OA | 63 | 91 | 81 | 0 | 18 | 29 |
| TBB | - OA | 91 | **100** | **100** | 91 | **100** | 41 |
|  | + OA | 98 | **100** | **100** | **100** | **100** | 91 |
| Saracatinib | - OA | 41 | 81 | 52 | 0 | 0 | 5 |
|  | + OA | **100** | **100** | **100** | 0 | 84 | 20 |
| A-107 | - OA | 92 | 79 | 65 | 82 | **100** | 80 |
|  | + OA | 36 | 70 | 85 | 21 | **100** | 55 |
| AR | - OA | **100** | 89 | **100** | **100** | **100** | **100** |
|  | + OA | 89 | **100** | **100** | **100** | **100** | **100** |
| CsA | - OA | 90 | 91 | 89 | 95 | 92 | 95 |
|  | + OA | 33 | 26 | 0 | 86 | 11 | 28 |

In contrast to N2a cells, LiCl caused considerable reduction of basal and OA-induced tau phosphorylation (monomeric p-tau, 67 kDa) in CTX culture at CP13 (-OA: 94%, +OA: 50%), RZ3 (-OA: 89%, +OA: 100%), PHF-1 (-OA: 98%, +OA: 100%), AT8 (-OA: 100%, +OA: 81%), AT270 (-OA: 100%, +OA: 100%) and total tau DA31 (-OA:93%, +OA:96%) (Figure 4a, 4b, lane 4 and Figure 5a 5b, lane 5, Table 4). AR also abolished the 67 kDa band with basal and OA-induced tau hyperphosphorylation (Figure 4a, 4b, lane 10 and Figure 5a, 5b, lane 11, Table 4). With total tau DA31 (a.a. 150-190), LiCl and AR completely reduced the 67 kDa (monomeric p-tau) and substantially increased the 63 kDa band (non-phospho tau). Treating CTX cells with A107 also showed a substantial inhibition of 67 kDa (-OA and +OA) band with CP13 (-OA: 92%, +OA: 36%), RZ3 (-OA: 79%, +OA: 70%), PHF-1 (-OA: 65%, +OA: 85%), AT8 (-OA: 82%, +OA: 21%), AT270 (-OA: 100%, +OA: 100%), and total tau DA31 (-OA: 80%, +OA: 55%), compared to OA treatment alone (Figure 4a, 4b, lane 9 and Figure 5a 5b, lane 10, Table 4). As for Roscovitine treatment, in contrast to N2a neuronal treatment, the 67 kDa band was reduced considerably at CP13 (-OA: 85%, +OA: 63%), RZ3 (-OA: 42%, +OA: 91%), PHF-1 (-OA: 63%, +OA: 81%), and total tau DA31 (-OA, +OA: ∼30%). However, Roscovitine did not show a statistically significant effect on OA-induced tau phosphorylation at AT8 and AT270 (Figure 4a, 4b, lane 6 and Figure 5a 5b, lane 7, Table 4).

On the other hand, CsA caused a molecular weight shift in the electrophoretic mobility of the 67 kDa to 63 kDa at the sites CP13 (pSer202), RZ3 (pThr231), and DA31 (a.a.102-145), presumably accounting for the dephosphorylation of tau (**Supplementary Figure 3**). In reference to the 67 kDa band, CsA had dramatic inhibition on basal tau phosphorylation at: CP13 (90%), RZ3 (91%), PHF-1 (89%), AT8 (95%), AT270 (92%) and total tau DA31 (67 kDa, 95%) (**Supplementary Figure 3a, b**, Table 4). With OA treatment, CsA also showed a considerable immunoreactivity reduction of 67 kDa band at the epitopes: CP13 (33%), AT8 (86%) and total tau DA31 (28%). CsA had no effect on 67 kDa band at PHF-1, AT270, and RZ3 compared to OA treatment alone (**Supplementary Figure 3a, b**, Table 4). Minor oligomeric bands were observed at 240 kDa with PHF-1 antibody in OA treated samples. Based on the αII-spectrin blot, CTX cultures demonstrated intact spectrin (240 kDa) and the absence of any SBDPs, suggesting a healthy metabolism under the experimental conditions.

Furthermore, Saracatinib caused considerable reduction of basal and OA-induced tau hyperphosphorylation at: CP13 (-OA: 41%, +OA: 100%), RZ3 (-OA: 81%, +OA: 100%), PHF-1 (-OA: 52%, +OA: 100%), AT270 (-OA: 0%, +OA: 84%) and total tau DA31 (-OA: 5%, +OA: 20%). Saracatinib did not show any significant effect at AT8 phospho-tau epitope (pSer202/pThr205 sites) (Figure 4a, 4b, lane 8 and Figure 5a, 5b, lane 9; Table 4).

Treatment with K252a caused substantial inhibition of 67 kDa band, with basal and OA-induced treatments at CP13 (-OA: 35%, +OA: 41%), RZ3 (-OA: 45%, +OA: 37%), PHF-1 (-OA: 61%, +OA: 63%), and total tau DA31 (-OA: 41%, +OA: 45%) (Figure 4a, 4b, lane 2, and Figure 5a, 5b, lane 3; Table 4). K252a did not show any statistically significant inhibition at AT8 and AT270 with both basal and OA-induced tau hyperphosphorylation (Figure 4a, 4b, lane 2; 5a, 5b, lane 3; Table 4). Cultures treated with STS showed considerable reduction of basal and OA-induced tau phosphorylation at CP13 (-OA: 86%, +OA: 63%), RZ3 (-OA: 83%, +OA: 81%), PHF-1 (-OA: 55%, +OA: 89%), AT8 (-OA: 88%, +OA: 12%), AT270 (-OA: 100%, +OA: 100%), and total tau DA31 (-OA: 41%, +OA: 45%) (Figure 4a, 4b, lane 3a and Figure 5a, 5b, lane 4; Table 4).

Unexpectedly, EGTA caused an adverse effect in CTX culture by further enhancing physiological p-tau and OA-induced tau hyperphosphorylation at CP13 (-OA: −51%,+OA:-12%), RZ3 (-OA:- 63%,+OA:-22%), PHF-1 (-OA:-22%,+OA:-13%), AT8 (-OA:-64%,+OA:-5%), AT270 (-OA:-68%,+OA:-63%), and total tau DA31 (-OA:-73%,+OA:-69%)(Figure 4a, 4b, lane 5 and Figure 5a, 5b, lane 6; Table 4). β-actin protein levels remained even in all experimental conditions. When samples were probed with αII-spectrin antibody, with the calpain and caspase-3 inhibitors added (S and Z), SBDP150/145 and SBDP120 bands were statistically non-significant compared to control, in both basal and OA-induced tau hyperphosphorylation, suggesting a healthy neuronal culture.

Taken all together, treatments with CKII inhibitor TBB, GSK3 inhibitors LiCl and AR, and Src/Fyn Kinase inhibitor Saracatinib showed robust inhibition leading to different reduced basal and OA-induced tau phosphorylation profiles demonstrating the specificity of inhibitors tested in our tauopathy cell-based models. Thus, the kinase inhibitors studied provide targets to reduce or prevent tau hyperphosphorylation and aggregation in tauopathies.

## Discussion

In the present study, OA was used to induce tau hyperphosphorylation and oligomerization in mouse neuroblastoma N2a culture and rat primary cerebrocortical neuronal cultures (CTX) to screen for various tau kinase inhibitors as potential drug candidates. The N2a neuronal cultures have been widely used to study mechanisms of neurodegeneration because they are a homogenous culture system that is convenient to handle and can multiply quickly to produce a tremendous amount of neuron precursor cells (72). However, the primary cultures were implemented in this study as they represent a healthier form of cortical neurons as opposed to cell lines which are cancerous, in a sense that gene expression in primary cortical culture could represent and mimic the actual *in vivo* expression. Additionally, primary culture has the advantage in portraying the complexity of the central nervous system by better translating into *in vivo* models used for screening pharmaceutical drug candidate’s compounds (73). It has been reported that OA results in robust tau hyperphosphorylation at multiple pathological epitopes in animal and cell culture studies (38, 40, 41, 74–76).

OA treatment was used to induce tau hyperphosphorylation and oligomerization at various phospho-tau epitopes in N2a cell culture as a tauopathy model. OA caused down-regulation of protein phosphatase and showed the appearance of oligomeric forms of p-tau species (110 kDa, 170 kDa, and 240 kDa) immunoreactive to p-tau-specific antibodies (pSer202, pSer396/404) and total anti-tau (a.a. 102-142). Although the exact mechanism of tauopathy-induced disorders is not yet elucidated, the immunostaining of autopsy brains with anti-p-tau antibodies, including AT8 (pSer202/pThr205), and PHF-1 (pSer396/pSer404) are utilized as a diagnostic method of AD and tauopathy (77). Thus, in our study, the increase in tau phosphorylation detected was identified at these sites as a representation of a tauopathy model.

Moreover, among all the phospho-tau epitopes studied here, Thr231 epitope is thought to be associated in the initiation of tau hyperphosphorylation in tauopathies while other epitopes such as Thr181, Ser202/Thr205, and Ser396/Ser404 are phosphorylated far ahead during the tauopathy process and the progression of the disease (78). The phosphorylation sites Thr231 and Thr181 have been proposed as biomarkers in AD while Ser202/Thr205 are used to determine the stage of AD progression (79–81). These phosphorylation sites were selected in our study to associate the effectiveness of protein kinase inhibitors with tauopathy-relevant phosphorylation sites.

It is widely established that PP2A is the primary enzyme responsible for dephosphorylation of tau protein throughout the brain, controlling all tau phosphorylation sites. PP2A activity is decreased in AD and TBI brains (12, 82). Therefore, the OA-induced inhibition of PP2A is a highly relevant model to study various tau protein kinase inhibitors as modulators of tau hyperphosphorylation and oligomerization targeting tau pathology (Figure 6, Table 2).

**Figure 6.**
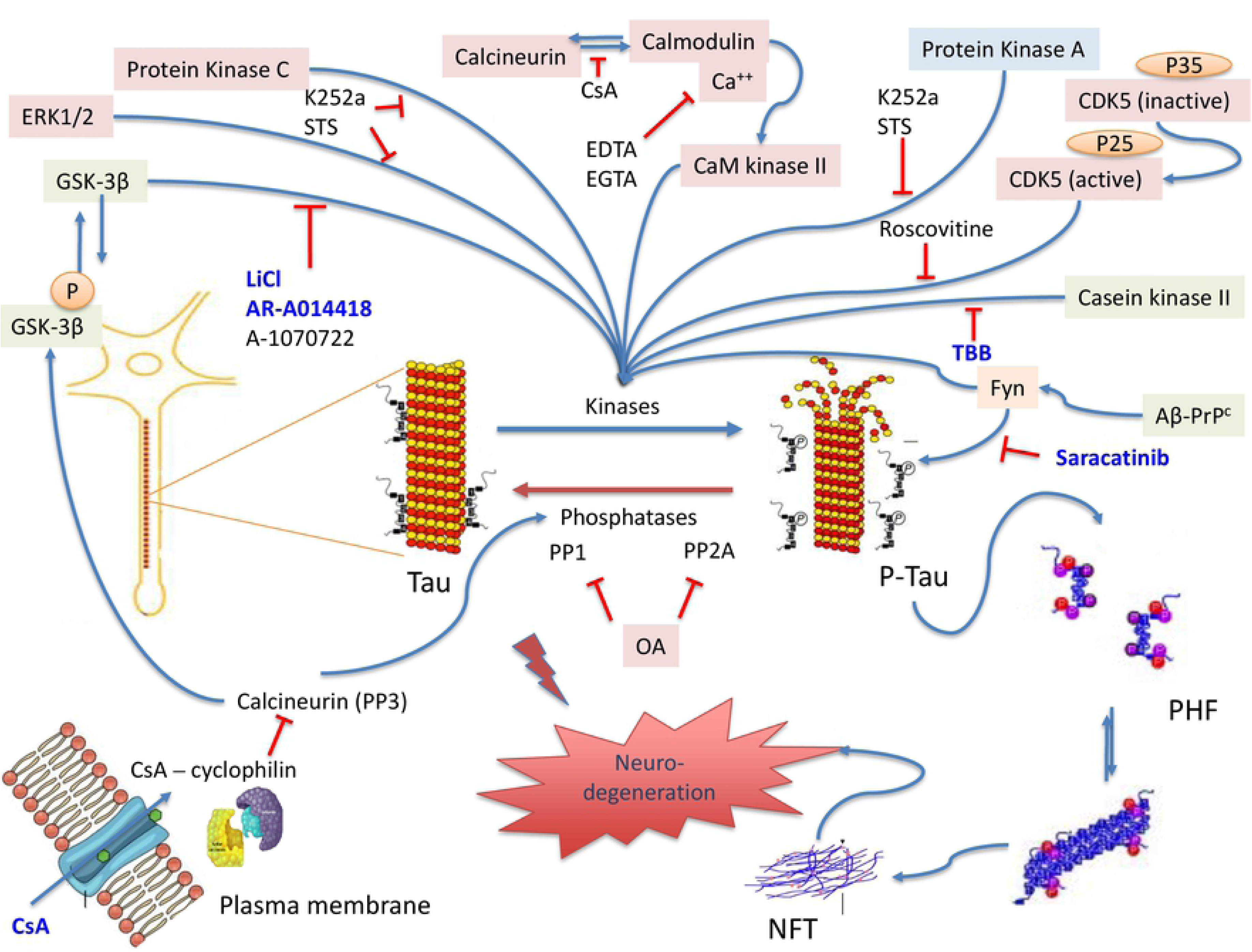
The tauopathy-model and a proposed mechanism for various protein kinase inhibitors intervention. Dephosphorylated tau protein binds the microtubules to maintain it in the polymerized state. Phosphorylation of tau protein by a host of different kinases causes tau to dissociate from the microtubules. Dissociation of tau causes the microtubules to depolymerize. Specific phosphatases dephosphorylate tau allowing the microtubule to re-polymerize again, a physiological process that provides structure and shape to the cytoskeleton of neurons. In tauopathies, imbalances between kinases and phosphatases functions lead to tau hyperphosphorylation at particular pathological sites and a higher tendency to dissociate from the microtubules producing soluble tau aggregates and insoluble paired helical filaments (PHF), that could combine to form neurofibrillary tangles (NFT). NFT is known to be the toxic species in AD and CTE, including other tauopathy diseases and little is known about their active mechanism of neurodegeneration. OA inhibits the function of crucial tau phosphatases, (PP1 and PP2A) leading to activation of tau kinases and tau hyperphosphorylation initiating the pathological processes of tauopathies. One pharmaceutical approach to reverse the mechanism of tauopathies is kinase inhibition. The protein kinase inhibitors selected in this study are indicated on this figure. The inhibitors highlighted in blue are ones that showed a promising effect on our OA-induced cell-based tauopathy model. Microsoft PowerPoint was used to create the artwork.

In contrast to N2a cell culture, the monomeric form of tau (ranging from 63 kDa – 67 kDa) was only observed in CTX culture. OA might be able to cause tau oligomerization inducing phosphorylation at different sites on tau depending upon the cell line. One possible reason for such effect would be that our separation of tau protein by SDS-PAGE was carried out under reducing conditions (Dithiothreitol (DTT) and β-ME) that could have the ability to minimize tau oligomers to the monomeric form, specifically in CTX primary culture. Our CTX serum-free neurobasal media contained antioxidants such as glutathione. Thus, the addition of these antioxidants may have blocked the process of tau oligomerization from occurring.

In the N2a and CTX cell culture, TBB (CKII inhibitor) surprisingly provided the most profound reversal of tau phosphorylation and oligomerization at the epitopes pSer202, (CP13), pSer396/pSer404 (PHF-1), pSer202/pThr205 (AT8), pThr181 (AT8), and pThr231 (RZ3). TBB is a selective, cell-permeable, ATP/GTP-competitive inhibitor of casein kinase II (CKII) (IC_50_=900 nM for rat liver) (83). It has been shown that CKII function is aberrant in AD, and its alteration precedes hyperphosphorylated tau accumulation in NFT formation (18). Moreover, it has been reported that CKII can phosphorylate tau purified from human brain and neuroblastoma cell line (18, 83–85). A study has shown that CKII phosphorylates SET, a potent PP2A inhibitor, inducing tau hyperphosphorylation in neurons and animal models, while inhibition of CKII by TBB eliminated this event (86). Thus, inhibition of CKII by TBB might provide a pharmacological interference for treating tauopathy-related disorders.

Since GSK-3 is a well-known kinase that can phosphorylate tau *in vitro* and *in vivo* and has been proposed as a target for pharmacological intervention (87, 88), three GSK-3 small molecule kinase inhibitors (LiCl, AR, and A-107) were selected to be assessed on OA-induced tauopathy, cell-based model. AR provided robust suppression of tau hyperphosphorylation in CTX culture at all tau epitopes tested (Figure 4a, 4b, and Figure 5a, 5b; and Table 4) and was less effective in N2a cells (Figure 2a, 2b, and Table 3). It was also observed that the effect of AR is more prominent compared to another GSK3 inhibitor, A-107 in CTX primary culture. This effect could be attributed, in part, to the high selectivity and specificity of AR to GSK3β (89) compared to A-107. A-107 display selectivity for both GSK3α and GSK3β (K_i_= 0.6 nM for both) (90) thereby might dilute the effect of inhibition of GSK3β, which is regarded as the critical kinase in AD (88).

Similarly, a study has shown that hypothermia-induced tau hyperphosphorylation was reduced with AR treatment in human neuroblastoma SH-SY5Y 3R-Tau (76). In another study, AR protected N2a cell culture against apoptosis by inhibition of the phosphatidylinositol-3 kinase/protein kinase B pathway and showed neuroprotective properties against neurotoxicity caused by the β-amyloid peptide in hippocampal slices (91). The lack of AR effect on N2a culture might be attributed to differences in cellular mechanisms from CTX culture, mediating OA-induced tau phosphorylation at multiple levels and different sites.

LiCl is well-known to inhibit GSK3 and other kinases (76). In CTX culture, LiCl caused dramatic inhibition of basal and OA-induced tau hyperphosphorylation at all tested tau epitopes. Consistent with previous reports, LiCl was shown to reduce tau phosphorylation in cultured cells, Ex-vivo rat brain slices, and rat brains at different AD-related tau epitopes (58, 76, 89, 92–94). Unexpectedly, LiCl showed an opposite impact on N2a culture by increasing OA-induced tau hyperphosphorylation and oligomerization at multiple tested tau epitopes. To the best of our knowledge, this effect is reported herein for the first time in cell culture. However, there are biological targets for LiCl that might have resulted in an adverse event. For instance, one hypothesis states that LiCl is a competitive inhibitor of GSK-3 to Mg^2+^, but not competitive to the substrate or ATP. Another theory proposes that LiCl causes potassium deprivation (95).

The use of CDK5 inhibitor Roscovitine in CTX culture substantially reduced basal and OA-induced tau hyperphosphorylation at CP13 (pSer202), RZ3 (pT231, PHF-1(pSer396/pSer404) and AT270 (pThr181). Roscovitine reduced basal phosphorylation at AT8 (pSer202/pThr205) but did not affect the OA-induced tau hyperphosphorylation, reflecting its specificity and the selectivity to our cell models. Similarly, several recent studies revealed that inhibiting CDK5 with Roscovitine had neuroprotective properties against neurodegenerative conditions caused by decreasing tau phosphorylation (66, 76, 96). Like LiCl, Roscovitine resulted in opposite effects in the N2a cells by increasing phosphorylation at CP13 (pSer202) and PHF-1 (pSer396/pSer404).

Another protein kinase that has recently received consideration as a pharmaceutical target is the tyrosine kinase Fyn, which has been linked with the amyloid pathway and tau phosphorylation through the N-terminal domain in dendrites (16). Saracatinib (also known as AZD0530) is a small molecular inhibitor that has high potency for Src and Fyn kinases (16, 19–21). Fyn can physically associate with tau and phosphorylate residues by interacting through its SH3 domain with SH3-binding domains in tau (Figure 6) (97). In our experiments, Saracatinib reduced both basal and OA-induced tau hyperphosphorylation (67 kDa) in N2a and CTX primary cultures at the epitopes: CP13 (pSer202), RZ3 (pThr231), PHF-1 (pSer396/pSer404) and AT270 (pThr181). Saracatinib did not affect the pSer202/pThr205 (AT8) site, suggesting that Fyn does not phosphorylate Thr205 residue in our experimental tauopathy model.

Cyclosporine (CsA) or FK506 is an 11 amino acid cyclic non-ribosomal peptide used as an immunosuppressant. CsA is known to induce neuroprotective properties through inhibiting specifically enzyme activity by binding to cyclophilin, forming a complex that inhibits calcineurin (PP3) (98) (Figure 6, Table 2). Several findings have shown that calcineurin inhibition increases tau hyperphosphorylation, and cells treated with CsA could induce the process (99, 100). In the present study, it was found that treatment with CsA alone did not result in any significant increase in tau levels or tau phosphorylation, which lies in agreement with a study done similarly (101), and reported complete inhibition of OA-induced tau hyperphosphorylation and oligomerization in N2a cells at the examined tau epitopes. In CTX culture, CsA produced a lower but still considerable reduction of OA-induced tau phosphorylation compared to N2a neuronal culture. These data suggest that PP2A is the main enzyme that regulates tau dephosphorylation in our culture system rather than PP3 at the tested sites. Moreover, we propose that CsA inhibits PP3 by blocking its binding to the calcium-dependent calmodulin, required for CaMKII to be active, thereby decreasing tau hyperphosphorylation (Figure 6, Table 2).

## Conclusions

In this study, OA was used to induce tauopathy in neuroblastoma and differentiated neuronal culture and screen for various pharmaceutical drug candidates. We provided a side-by-side comparison of possible drug candidates that are well described in respect to tauopathies such as Alzheimer’s (Saracatinib, LiCl, AR) as well as other prospects which have been minimally studied in application to potential therapies (TBB and CsA). TBB and CsA warrant further test design involving an animal model of tauopathy. This is particularly important as recent studies implicate pre-fibrillar hyperphosphorylated tau as the toxic species in AD, CTE, and other neurodegenerative diseases, therefore, re-establishing the interest in tau kinase inhibitors development at putative neurotherapies, which could translate into human clinical trials.

## Limitation and Future Directions

With the use of peptidomic, it is evident that unique peptides are the main causes of tauopathies; we hypothesized that after using OA induced tau hyperphosphorylation as a tauopathy model, different drugs could be tested as neurotherapeutic targets for tauopathies-related disorders. In one study, different Kinase inhibitors were tested on samples, to model possible therapeutic drugs. TBB is a novel CKII inhibitor that has not been used for tauopathies and can be an example for other possible drugs.

The idea of targeting specific biomarkers, after Peptidomic screening helped identified potential kinase inhibitor candidates (AR, TBB, CsA, and Saracatinib) that warrant further test design. The study would like to further test this protocol on 3R vs 4R human tau cell cultures. Experiments on CSF and blood samples are being run to make up for the high based evidence on rodent cultures.

Some limitations to the study include the idea that these compound dilutions are arbitrary or based on references using different cell systems. Treatment is often in the high micromolar range where compounds acts on more than their primary target. The use of serial dilution and the generation of an EC50 would have been more appropriate.

Furthermore, the technic used is entirely based on western blotting which prohibits its potential use in higher throughput screens. These systems are highly specific, but a wider variety of auto-antibody assays, would have provided more accurate results.

Finally, one constraint to this study remain from a lack of interpretation on why some compounds respond in basal conditions while other respond in OA-treated conditions. The discrepancy in response between the N2a cell line and CTX also cautions the interpretation of the screen. Having a wider variety of samples from different parts of the brain, could have potentially given more precise evidence toward the study.

## List of Abbreviations

(A-1070722): 1-(7-methoxyquinolin-4-yl)-3-(6-(trifluoromethyl) pyridin-2-yl
AD: Alzheimer’s disease
AR-A014418: N-(4-methoxybenzyl)-N’-(5-nitro-1,3-thiazol-2-yl) urea
AZD0530: Saracatinib
BCIT/NBT: 5-bromo-4-chloro-3-indolyl phosphate
CaMKII: Ca2+/calmodulin-dependent protein kinase II
CDK5: Cyclin-dependent kinase 5
CK2: Casein kinase
CKII: Casein kinase II
CsA: Cyclosporin A
CTE: Chronic traumatic encephalopathy
CTX: Neuronal cortical cell cultures
DTT: Dithiothreitol
DYRK1A/2: Dual-specificity tyrosine kinase phosphorylation and regulated kinase-1A/2
EGTA: Ethylene glycol-bis (β-aminoethyl ether)-N,N,N’,N’-tetraacetic acid
GSK-3: Glycogen synthase kinase 3 inhibitors
HMW: High Molecular Weight
K252a: Staurosporine analog, non-selective cell permeable Protein Kinase Inhibitor; (9S,10R,12R)-2,3,9,10,11,12-Hexahydro-10-hydroxy-9-methyl-1-oxo-9,12-epoxy-1H-diindolo[1,2,3-fg:3’,2’,1’-kl]pyrrolo[3,4-i][1,6]benzodiazocine-10-carboxylic acid methyl ester
LiCl: Lithium Chloride
LMW-MT: Low molecular weight monomeric tau
MLCK: Myosin light-chain kinase
N2A: Neuroblastoma cell line
NFT: Neurofibrillary tangles
NMDAR: N-methyl-D-aspartate receptors
OA: Okadaic acid
PP1: Protein Phosphatase 1
PP2A: Protein Phosphatase 2A
Roscovitine: (2R)-2-1-butanol
Saracatinib: N-(5-chloro-1,3-benzodioxol-4-yl)-7-[2-(4-methylpiperazin-1-yl)ethoxy]-5-(tetrahydro-2H-pyran-4-yloxy)quinazolin-4-amine
SBDP: Spectrin breakdown products
SNJ-1945: Amphipathic ketoamide
STS: Staurosporine; (5S,6R,7R,9R)-6-methoxy-5-methyl-7-(methylamino)-6,7,8,9,15,16-hexahydro-17-oxa-4b,9a,15-triaza-5,9-methanodibenzo[b,h]cyclonona[jkl]cyclopenta[e]-as-indacen-14(5h)-one
TBB: 4,5,6,7-tetrabromobenzotriazole
Z-DCB: Caspase-3; Z-Asp-2,6-Dichlorobenzoyloxymethyl Ketone

## Declarations

### Availability of data and material

- All data generated or analyzed during this study are included in this article (and its additional files).

### Competing interests

- The authors declare that they have no conflict of interest.

### Funding

- This work was supported by funding from the University of Florida Departments of Emergency Medicine and Psychiatry.

## Authors’ contributions

Conceived and designed the experiments: HY, KW. Performed the experiments: HY, IT, GA, MK. Analyzed the Data: HY, KW. Contributed reagents/material/analysis tools: ZY, FL, and PD. Performed revision experiments: HY and KW. Wrote the manuscript: HY, MK Reviewed the manuscript: RY, KW, and FK.

## Acknowledgment

We would like to thank Dr. Peter Davies (Albert Einstein College of Medicine, Bronx, NY, USA) for the kind gift of monoclonal tau antibodies.

---

**Supplementary Figure 1: Effect of cyclosporin A on OA-induced tau hyperphosphorylation in mouse N2a cells.**

The same experimental design mentioned in Figure 2 was used to test CsA in N2a cell culture. Twenty micrograms of protein extract were used for the analysis of tau. Calpain and caspase-3 inhibitors (S+Z) were added to all experimental conditions, including the control samples. CsA is known to inhibit the phosphatase activity of calcineurin (PP3). In the presented experiment, it is used to assess its kinase inhibition potential on the monomeric and oligomeric p-tau induced by OA. (a). Immunoblots of N2a neuronal culture protein extracts showing antibodies directed against major tau phosphorylation sites. Two additional p-tau antibodies were used (AT270 and RZ3) to assess the phosphorylation sites at pThr181 and pThr231, respectively. RZ3 and AT270 detected distinctive monomeric p-tau bands at 48 kDa, and 55 kDa, respectively. Total tau levels were probed using DA9 (a.a. 102-145) in N2a cells. Blue colored labels correspond to monomeric or oligomeric p-tau species. Immunoblots were probed with αII-spectrin antibody to monitor calpain and caspase-3 mediated proteolysis. (b). Immunoblots quantification of N2a. The ratio of phosphorylation epitopes levels over β-actin levels ± SD are represented as a percentage of control. n=3 per condition. For multiple comparisons, one-way ANOVA followed by the Bonferroni’s post-hoc test was performed. *p<0.05, **p<0.01, ***p<0.001, ****p<0.0001, ns: non-significant.

**Supplementary Figure 2. Effect of additional two GSK-3 protein kinase inhibitors on OA-induced tau hyperphosphorylation and oligomerization in N2a cells (with cell-death linked protease inhibitors (calpain/caspase inhibitors).**

A continuation of Figure 2 experiment is presented to include two other potent GSK-3 kinase inhibitors, AR and A-107. The detailed experimental treatments are as described in materials and methods. (a). Immunoblots of N2a cells extracted protein using p-tau antibodies (CP13 and PHF-1), total tau (DA9), and αII-Spectrin. αII-Spectrin was probed to assess cell apoptosis monitored SBDP150/145 kDa and SBDP120 kDa. Kinase inhibition of phosphorylation and oligomerization was monitored by evaluating the levels of p-tau antibodies and total tau (blue arrows) and non-phospho tau (black arrows). For all conditions, S+Z were added for 1h before the treatments. (b). Immunoblots quantification and statistical analysis. All data are normalized to β-actin and are expressed as a percentage of control. Data are presented as ± SEM for n=3. Statistical analysis was performed with one-way ANOVA. For multiple comparisons, one-way ANOVA followed by the Bonferroni’s post-hoc test was performed. *p<0.05, **p<0.01, ***p<0.001, ****p<0.0001 and ns: non-significant.

**Supplementary Figure 3. Cyclosporin A inhibits physiological and OA-induced Tau hyperphosphorylation in rat primary cerebrocortical neuronal culture.**

The experimental procedures were followed as described in Figure 4 and Figure 5 legends. Primary neuronal cultures (CTX) were fully differentiated and had healthy neurites when examined under the microscope. All wells were pretreated with S+Z for 1h. For conditions that did not include OA, cultures were treated with CsA for 6h. For OA-induced conditions, OA was added for 24h followed by CsA for 6h. A reverse time course was followed, and all experimental conditions were collected and analyzed at the same time. Twenty micrograms of CTX culture extracts were run on SDS-PAGE followed by western blotting. (a). Immunoblots of CTX culture protein extracts. CTX culture using antibodies directed against major tau phosphorylation sites including: CP13 (pSer202), PHF-1 (pSer396/pSer404), RZ3 (pThr231), AT8 (pSer202/pThr205), AT270 (pThr181). Total tau levels were probed using DA31 (a.a. 102-145). The 67 kDa assigned as monomeric p-tau band and the 63 kDa band was assigned as monomeric non-phospho tau at the different studied epitopes. (b). Immunoblots quantification. The ratio of phosphorylation epitopes levels over β-actin levels ± SD are represented as a percentage of control. n=3 per condition. For multiple comparisons, one-way ANOVA followed by the Bonferroni’s post-hoc test was performed. *p<0.05, **p<0.01, ***p<0.001, ****p<0.0001, ns: non-significant.

**Supplementary Figure 4. Full scanned immunoblots for Figure 1a.**

**Supplementary Figure 5. Full scanned immunoblots for Figure 2a.**

**Supplementary Figure 6. Full scanned immunoblots for Figure 3a.**

**Supplementary Figure 7. Full scanned immunoblots for Figure 4a.**

**Supplementary Figure 8. Full scanned immunoblots for Figure 5a.**

## References

1. Sparks P, Lawrence T, Hinze S. Neuroimaging in the Diagnosis of Chronic Traumatic Encephalopathy: A Systematic Review. Clin J Sport Med. 2017.

2. Panza F, Imbimbo BP, Lozupone M, Greco A, Seripa D, Logroscino G, et al. Disease-modifying therapies for tauopathies: agents in the pipeline. Expert Rev Neurother. 2019.

3. Mohamed AZ, Cumming P, Gotz J, Nasrallah F, Department of Defense Alzheimer’s Disease Neuroimaging I. Tauopathy in veterans with long-term posttraumatic stress disorder and traumatic brain injury. Eur J Nucl Med Mol Imaging. 2019;46(5):1139–51.

4. Wang L, Benzinger TL, Su Y, Christensen J, Friedrichsen K, Aldea P, et al. Evaluation of Tau Imaging in Staging Alzheimer Disease and Revealing Interactions Between beta-Amyloid and Tauopathy. JAMA Neurol. 2016;73(9):1070–7.

5. Besser LM, Mock C, Teylan MA, Hassenstab J, Kukull WA, Crary JF. Differences in Cognitive Impairment in Primary Age-Related Tauopathy Versus Alzheimer Disease. J Neuropathol Exp Neurol. 2019.

6. Pir GJ, Choudhary B, Mandelkow E. models of tauopathy. FASEB J. 2017;31(12):5137–48.

7. Perrine K, Helcer J, Tsiouris AJ, Pisapia DJ, Stieg P. The Current Status of Research on Chronic Traumatic Encephalopathy. World Neurosurg. 2017;102:533–44.

8. Kovacs GG. Tauopathies. Handb Clin Neurol. 2017;145:355–68.

9. Neve RL, Harris P, Kosik KS, Kurnit DM, Donlon TA. Identification of cDNA clones for the human microtubule-associated protein tau and chromosomal localization of the genes for tau and microtubule-associated protein 2. Brain Res. 1986;387(3):271–80.

10. Avila J, Jiménez JS, Sayas CL, Bolós M, Zabala JC, Rivas G, et al. Tau Structures. Front Aging Neurosci. 2016;8:262.

11. Lee G, Leugers CJ. Tau and Tauopathies. Prog Mol Biol Transl Sci. 2012;107:263–93.

12. Theendakara V, Bredesen DE, Rao RV. Downregulation of protein phosphatase 2A by apolipoprotein E: Implications for Alzheimer’s disease. Mol Cell Neurosci. 2017;83:83–91.

13. Stoothoff WH, Johnson GV. Tau phosphorylation: physiological and pathological consequences. Biochim Biophys Acta. 2005;1739(2-3):280–97.

14. Ni R, Ji B, Ono M, Sahara N, Zhang MR, Aoki I, et al. Comparative in-vitro and in-vivo quantifications of pathological tau deposits and their association with neurodegeneration in tauopathy mouse models. J Nucl Med. 2018.

15. Sahara N, Shimojo M, Ono M, Takuwa H, Febo M, Higuchi M, et al. Tau Imaging for a Diagnostic Platform of Tauopathy Using the rTg4510 Mouse Line. Front Neurol. 2017;8:663.

16. Nygaard HB. Targeting Fyn Kinase in Alzheimer’s Disease. Biol Psychiatry. 2018;83(4):369–76.

17. Martin L, Latypova X, Wilson CM, Magnaudeix A, Perrin ML, Yardin C, et al. Tau protein kinases: involvement in Alzheimer’s disease. Ageing Res Rev. 2013;12(1):289–309.

18. Rosenberger AF, Morrema TH, Gerritsen WH, van Haastert ES, Snkhchyan H, Hilhorst R, et al. Increased occurrence of protein kinase CK2 in astrocytes in Alzheimer’s disease pathology. J Neuroinflammation. 2016;13:4.

19. Li C, Götz J. Somatodendritic accumulation of Tau in Alzheimer’s disease is promoted by Fyn-mediated local protein translation. EMBO J. 2017;36(21):3120–38.

20. Liu W, Zhao J, Lu G. miR-106b inhibits tau phosphorylation at Tyr18 by targeting Fyn in a model of Alzheimer’s disease. Biochem Biophys Res Commun. 2016;478(2):852–7.

21. Kaufman AC, Salazar SV, Haas LT, Yang J, Kostylev MA, Jeng AT, et al. Fyn inhibition rescues established memory and synapse loss in Alzheimer mice. Ann Neurol. 2015;77(6):953–71.

22. Dolan PJ, Johnson GV. The role of tau kinases in Alzheimer’s disease. Curr Opin Drug Discov Devel. 2010;13(5):595–603.

23. Bennett PC, Zhao W, Ng KT. Concentration-dependent effects of protein phosphatase (PP) inhibitors implicate PP1 and PP2A in different stages of memory formation. Neurobiol Learn Mem. 2001;75(1):91–110.

24. Ando K, Maruko-Otake A, Ohtake Y, Hayashishita M, Sekiya M, Iijima KM. Stabilization of Microtubule-Unbound Tau via Tau Phosphorylation at Ser262/356 by Par-1/MARK Contributes to Augmentation of AD-Related Phosphorylation and Abeta42-Induced Tau Toxicity. PLoS Genet. 2016;12(3):e1005917.

25. Gerson JE, Farmer KM, Henson N, Castillo-Carranza DL, Carretero Murillo M, Sengupta U, et al. Tau oligomers mediate alpha-synuclein toxicity and can be targeted by immunotherapy. Mol Neurodegener. 2018;13(1):13.

26. Tan CC, Zhang XY, Tan L, Yu JT. Tauopathies: Mechanisms and Therapeutic Strategies. J Alzheimers Dis. 2018;61(2):487–508.

27. Wang KK, Yang Z, Zhu T, Shi Y, Rubenstein R, Tyndall JA, et al. An update on diagnostic and prognostic biomarkers for traumatic brain injury. Expert Rev Mol Diagn. 2018;18(2):165–80.

28. Aldag M, Armstrong RC, Bandak F, Bellgowan PSF, Bentley T, Biggerstaff S, et al. The Biological Basis of Chronic Traumatic Encephalopathy following Blast Injury: A Literature Review. J Neurotrauma. 2017;34(S1):S26–S43.

29. Panza F, Solfrizzi V, Seripa D, Imbimbo BP, Lozupone M, Santamato A, et al. Tau-based therapeutics for Alzheimer’s disease: active and passive immunotherapy. Immunotherapy. 2016;8(9):1119–34.

30. Tolosa E, Litvan I, Höglinger GU, Burn D, Lees A, Andrés MV, et al. A phase 2 trial of the GSK-3 inhibitor tideglusib in progressive supranuclear palsy. Mov Disord. 2014;29(4):470–8.

31. Georgievska B, Sandin J, Doherty J, Mörtberg A, Neelissen J, Andersson A, et al. AZD1080, a novel GSK3 inhibitor, rescues synaptic plasticity deficits in rodent brain and exhibits peripheral target engagement in humans. J Neurochem. 2013;125(3):446–56.

32. Ludolph AC, Kassubek J, Landwehrmeyer BG, Mandelkow E, Mandelkow EM, Burn DJ, et al. Tauopathies with parkinsonism: clinical spectrum, neuropathologic basis, biological markers, and treatment options. Eur J Neurol. 2009;16(3):297–309.

33. Medina M. An Overview on the Clinical Development of Tau-Based Therapeutics. Int J Mol Sci. 2018;19(4).

34. Tell V, Hilgeroth A. Recent developments of protein kinase inhibitors as potential AD therapeutics. Front Cell Neurosci. 2013;7:189.

35. Alonso AD, Di Clerico J, Li B, Corbo CP, Alaniz ME, Grundke-Iqbal I, et al. Phosphorylation of tau at Thr212, Thr231, and Ser262 combined causes neurodegeneration. J Biol Chem. 2010;285(40):30851–60.

36. Simic G, Babic Leko M, Wray S, Harrington C, Delalle I, Jovanov-Milosevic N, et al. Tau Protein Hyperphosphorylation and Aggregation in Alzheimer’s Disease and Other Tauopathies, and Possible Neuroprotective Strategies. Biomolecules. 2016;6(1):6.

37. Rabano A, Cuadros R, Merino-Serrais P, Rodal I, Benavides-Piccione R, Gomez E, et al. Protocols for Monitoring the Development of Tau Pathology in Alzheimer’s Disease. Methods Mol Biol. 2016;1303:143–60.

38. Kamat PK, Rai S, Swarnkar S, Shukla R, Ali S, Najmi AK, et al. Okadaic acid-induced Tau phosphorylation in rat brain: role of NMDA receptor. Neuroscience. 2013;238:97–113.

39. Ho YS, Yang X, Lau JC, Hung CH, Wuwongse S, Zhang Q, et al. Endoplasmic reticulum stress induces tau pathology and forms a vicious cycle: implication in Alzheimer’s disease pathogenesis. J Alzheimers Dis. 2012;28(4):839–54.

40. Jones NC, Nguyen T, Corcoran NM, Velakoulis D, Chen T, Grundy R, et al. Targeting hyperphosphorylated tau with sodium selenate suppresses seizures in rodent models. Neurobiol Dis. 2012;45(3):897–901.

41. Broetto N, Hansen F, Brolese G, Batassini C, Lirio F, Galland F, et al. Intracerebroventricular administration of okadaic acid induces hippocampal glucose uptake dysfunction and tau phosphorylation. Brain Res Bull. 2016;124:136–43.

42. Wang J, Tung YC, Wang Y, Li XT, Iqbal K, Grundke-Iqbal I. Hyperphosphorylation and accumulation of neurofilament proteins in Alzheimer disease brain and in okadaic acid-treated SY5Y cells. FEBS Lett. 2001;507(1):81–7.

43. Baker S, Gotz J. A local insult of okadaic acid in wild-type mice induces tau phosphorylation and protein aggregation in anatomically distinct brain regions. Acta neuropathologica communications. 2016;4:32.

44. Boban M, Babic Leko M, Miskic T, Hof PR, Simic G. Human neuroblastoma SH-SY5Y cells treated with okadaic acid express phosphorylated high molecular weight tau-immunoreactive protein species. J Neurosci Methods. 2019;319:60–8.

45. Shen XY, Luo T, Li S, Ting OY, He F, Xu J, et al. Quercetin inhibits okadaic acid-induced tau protein hyperphosphorylation through the Ca2+- calpain- p25- CDK5 pathway in HT22 cells. Int J Mol Med. 2018;41(2):1138–46.

46. Zhang Z, Simpkins JW. An okadaic acid-induced model of tauopathy and cognitive deficiency. Brain Res. 2010;1359:233–46.

47. Valdiglesias V, Prego-Faraldo MV, Pasaro E, Mendez J, Laffon B. Okadaic acid: more than a diarrheic toxin. Mar Drugs. 2013;11(11):4328–49.

48. Koumura A, Nonaka Y, Hyakkoku K, Oka T, Shimazawa M, Hozumi I, et al. A novel calpain inhibitor, ((1S)-1((((1S)-1-benzyl-3-cyclopropylamino-2,3-di-oxopropyl)amino)carbonyl)-3-methylbutyl) carbamic acid 5-methoxy-3-oxapentyl ester, protects neuronal cells from cerebral ischemia-induced damage in mice. Neuroscience. 2008;157(2):309–18.

49. Drognitz O, Obermaier R, Liu X, Neeff H, von Dobschuetz E, Hopt UT, et al. Effects of organ preservation, ischemia time and caspase inhibition on apoptosis and microcirculation in rat pancreas transplantation. Am J Transplant. 2004;4(7):1042–50.

50. Kobeissy FH, Liu MC, Yang Z, Zhang Z, Zheng W, Glushakova O, et al. Degradation of betaII-Spectrin Protein by Calpain-2 and Caspase-3 Under Neurotoxic and Traumatic Brain Injury Conditions. Mol Neurobiol. 2015;52(1):696–709.

51. Mondello S, Robicsek SA, Gabrielli A, Brophy GM, Papa L, Tepas J, et al. αII-Spectrin Breakdown Products (SBDPs): Diagnosis and Outcome in Severe Traumatic Brain Injury Patients. J Neurotrauma. 2010;27(7):1203–13.

52. Khélifa T, Beck WT. Induction of apoptosis by dexrazoxane (ICRF-187) through caspases in the absence of c-jun expression and c-Jun NH2-terminal kinase 1 (JNK1) activation in VM-26-resistant CEM cells. Biochem Pharmacol. 1999;58(8):1247–57.

53. Iimoto DS, Masliah E, DeTeresa R, Terry RD, Saitoh T. Aberrant casein kinase II in Alzheimer’s disease. Brain Res. 1990;507(2):273–80.

54. Toledo FD, Pérez LM, Basiglio CL, Ochoa JE, Sanchez Pozzi EJ, Roma MG. The Ca²⁺-calmodulin-Ca²⁺/calmodulin-dependent protein kinase II signaling pathway is involved in oxidative stress-induced mitochondrial permeability transition and apoptosis in isolated rat hepatocytes. Arch Toxicol. 2014;88(9):1695–709.

55. Iqbal K, Liu F, Gong CX, Grundke-Iqbal I. Tau in Alzheimer Disease and Related Tauopathies. Curr Alzheimer Res. 2010;7(8):656–64.

56. Qin N, Olcese R, Bransby M, Lin T, Birnbaumer L. Ca2+-induced inhibition of the cardiac Ca2+ channel depends on calmodulin. Proc Natl Acad Sci U S A. 1999;96(5):2435–8.

57. Song JS, Yang SD. Tau protein kinase I/GSK-3 beta/kinase FA in heparin phosphorylates tau on Ser199, Thr231, Ser235, Ser262, Ser369, and Ser400 sites phosphorylated in Alzheimer disease brain. J Protein Chem. 1995;14(2):95–105.

58. Noble W, Planel E, Zehr C, Olm V, Meyerson J, Suleman F, et al. Inhibition of glycogen synthase kinase-3 by lithium correlates with reduced tauopathy and degeneration in vivo. Proc Natl Acad Sci U S A. 2005;102(19):6990–5.

59. Prabhakaran J, Zanderigo F, Solingapuram Sai KK, Rubin-Falcone H, Jorgensen MJ, Kaplan JR, et al. Radiosynthesis and in Vivo Evaluation of [11C]A1070722, a High Affinity GSK-3 PET Tracer in Primate Brain. ACS Chem Neurosci. 2017;8(8):1697–703.

60. Gucalp A, Sparano JA, Caravelli J, Santamauro J, Patil S, Abbruzzi A, et al. Phase II trial of saracatinib (AZD0530), an oral SRC-inhibitor for the treatment of patients with hormone receptor-negative metastatic breast cancer. Clin Breast Cancer. 2011;11(5):306–11.

61. Tapley P, Lamballe F, Barbacid M. K252a is a selective inhibitor of the tyrosine protein kinase activity of the trk family of oncogenes and neurotrophin receptors. Oncogene. 1992;7(2):371–81.

62. Zimmermann A, Keller H. Effects of staurosporine, K 252a and other structurally related protein kinase inhibitors on shape and locomotion of Walker carcinosarcoma cells. Br J Cancer. 1992;66(6):1077–82.

63. Ruegg UT, Burgess GM. Staurosporine, K-252 and UCN-01: potent but nonspecific inhibitors of protein kinases. Trends in pharmacological sciences. 1989;10(6):218–20.

64. Liu MC, Kobeissy F, Zheng W, Zhang Z, Hayes RL, Wang KK. Dual vulnerability of tau to calpains and caspase-3 proteolysis under neurotoxic and neurodegenerative conditions. ASN Neuro. 2011;3(1):e00051.

65. Wang KK, Yang Z, Chiu A, Lin F, Rubenstein R. Examining the Neural and Astroglial Protective Effects of Cellular Prion Protein Expression and Cell Death Protease Inhibition in Mouse Cerebrocortical Mixed Cultures. Mol Neurobiol. 2016;53(7):4821–32.

66. Meijer L, Borgne A, Mulner O, Chong JP, Blow JJ, Inagaki N, et al. Biochemical and cellular effects of roscovitine, a potent and selective inhibitor of the cyclin-dependent kinases cdc2, cdk2 and cdk5. European journal of biochemistry. 1997;243(1-2):527–36.

67. Bhounsule AS, Bhatt LK, Prabhavalkar KS, Oza M. Cyclin dependent kinase 5: A novel avenue for Alzheimer’s disease. Brain Res Bull. 2017;132:28–38.

68. Engmann O, Giese KP. Crosstalk between Cdk5 and GSK3β: Implications for Alzheimer’s Disease. Frontiers in Molecular Neuroscience. 2009;2:2.

69. Sengupta A, Grundke-Iqbal I, Iqbal K. Regulation of phosphorylation of tau by protein kinases in rat brain. Neurochem Res. 2006;31(12):1473–80.

70. Davis DR, Brion JP, Couck AM, Gallo JM, Hanger DP, Ladhani K, et al. The phosphorylation state of the microtubule-associated protein tau as affected by glutamate, colchicine and beta-amyloid in primary rat cortical neuronal cultures. Biochem J. 1995;309 (Pt 3):941–9.

71. Sahara N, DeTure M, Ren Y, Ebrahim AS, Kang D, Knight J, et al. Characteristics of TBS-extractable hyperphosphorylated tau species: aggregation intermediates in rTg4510 mouse brain. J Alzheimers Dis. 2013;33(1):249–63.

72. Chen Y, Wang C, Hu M, Pan J, Chen J, Duan P, et al. Effects of ginkgolide A on okadaic acid-induced tau hyperphosphorylation and the PI3K-Akt signaling pathway in N2a cells. Planta Med. 2012;78(12):1337–41.

73. Schlachetzki JC, Saliba SW, Oliveira AC. Studying neurodegenerative diseases in culture models. Rev Bras Psiquiatr. 2013;35 Suppl 2:S92–100.

74. Zhao L, Xiao Y, Wang XL, Pei J, Guan ZZ. Original Research: Influence of okadaic acid on hyperphosphorylation of tau and nicotinic acetylcholine receptors in primary neurons. Exp Biol Med (Maywood). 2016;241(16):1825–33.

75. Hübinger G, Geis S, LeCorre S, Mühlbacher S, Gordon S, Fracasso RP, et al. Inhibition of PHF-like tau hyperphosphorylation in SH-SY5Y cells and rat brain slices by K252a. J Alzheimers Dis. 2008;13(3):281–94.

76. Bretteville A, Marcouiller F, Julien C, El Khoury NB, Petry FR, Poitras I, et al. Hypothermia-induced hyperphosphorylation: a new model to study tau kinase inhibitors. Sci Rep. 2012;2:480.

77. Kimura T, Sharma G, Ishiguro K, Hisanaga SI. Phospho-Tau Bar Code: Analysis of Phosphoisotypes of Tau and Its Application to Tauopathy. Front Neurosci. 2018;12:44.

78. Augustinack JC, Schneider A, Mandelkow EM, Hyman BT. Specific tau phosphorylation sites correlate with severity of neuronal cytopathology in Alzheimer’s disease. Acta Neuropathol. 2002;103(1):26–35.

79. Hampel H, Buerger K, Zinkowski R, Teipel SJ, Goernitz A, Andreasen N, et al. Measurement of phosphorylated tau epitopes in the differential diagnosis of Alzheimer disease: a comparative cerebrospinal fluid study. Arch Gen Psychiatry. 2004;61(1):95–102.

80. Hampel H, Goernitz A, Buerger K. Advances in the development of biomarkers for Alzheimer’s disease: from CSF total tau and Abeta(1-42) proteins to phosphorylated tau protein. Brain Res Bull. 2003;61(3):243–53.

81. Buerger K, Teipel SJ, Zinkowski R, Blennow K, Arai H, Engel R, et al. CSF tau protein phosphorylated at threonine 231 correlates with cognitive decline in MCI subjects. Neurology. 2002;59(4):627–9.

82. Lucke-Wold BP, Turner RC, Logsdon AF, Bailes JE, Huber JD, Rosen CL. Linking traumatic brain injury to chronic traumatic encephalopathy: identification of potential mechanisms leading to neurofibrillary tangle development. J Neurotrauma. 2014;31(13):1129–38.

83. Sarno S, Papinutto E, Franchin C, Bain J, Elliott M, Meggio F, et al. ATP site-directed inhibitors of protein kinase CK2: an update. Curr Top Med Chem. 2011;11(11):1340–51.

84. Avila J, Ulloa L, González J, Moreno F, Díaz-Nido J. Phosphorylation of microtubule-associated proteins by protein kinase CK2 in neuritogenesis. Cell Mol Biol Res. 1994;40(5-6):573–9.

85. Greenwood JA, Scott CW, Spreen RC, Caputo CB, Johnson GV. Casein kinase II preferentially phosphorylates human tau isoforms containing an amino-terminal insert. Identification of threonine 39 as the primary phosphate acceptor. J Biol Chem. 1994;269(6):4373–80.

86. Zhang Q, Xia Y, Wang Y, Shentu Y, Zeng K, Mahaman YAR, et al. CK2 Phosphorylating I2(PP2A)/SET Mediates Tau Pathology and Cognitive Impairment. Front Mol Neurosci. 2018;11:146.

87. Medina M, Garrido JJ, Wandosell FG. Modulation of GSK-3 as a Therapeutic Strategy on Tau Pathologies. Front Mol Neurosci. 2011;4:24.

88. Llorens-Martin M, Jurado J, Hernandez F, Avila J. GSK-3beta, a pivotal kinase in Alzheimer disease. Front Mol Neurosci. 2014;7:46.

89. Gould TD, Einat H, Bhat R, Manji HK. AR-A014418, a selective GSK-3 inhibitor, produces antidepressant-like effects in the forced swim test. Int J Neuropsychopharmacol. 2004;7(4):387–90.

90. Prabhakaran J, Zanderigo F, Sai KKS, Rubin-Falcone H, Jorgensen MJ, Kaplan JR, et al. Radiosynthesis and in Vivo Evaluation of [ACS Chem Neurosci. 2017;8(8):1697–703.

91. Bhat R, Xue Y, Berg S, Hellberg S, Ormo M, Nilsson Y, et al. Structural insights and biological effects of glycogen synthase kinase 3-specific inhibitor AR-A014418. J Biol Chem. 2003;278(46):45937–45.

92. Orena SJ, Torchia AJ, Garofalo RS. Inhibition of glycogen-synthase kinase 3 stimulates glycogen synthase and glucose transport by distinct mechanisms in 3T3-L1 adipocytes. J Biol Chem. 2000;275(21):15765–72.

93. Fu ZQ, Yang Y, Song J, Jiang Q, Lin ZC, Wang Q, et al. LiCl attenuates thapsigargin-induced tau hyperphosphorylation by inhibiting GSK-3beta in vivo and in vitro. J Alzheimers Dis. 2010;21(4):1107–17.

94. Lee S, Shea TB. Regulation of tau proteolysis by phosphatases. Brain Res. 2013;1495:30–6.

95. Kramer T, Schmidt B, Lo Monte F. Small-Molecule Inhibitors of GSK-3: Structural Insights and Their Application to Alzheimer’s Disease Models. Int J Alzheimers Dis. 2012;2012:381029.

96. Liu J, Yang J, Xu Y, Guo G, Cai L, Wu H, et al. Roscovitine, a CDK5 Inhibitor, Alleviates Sevoflurane-Induced Cognitive Dysfunction via Regulation Tau/GSK3beta and ERK/PPARgamma/CREB Signaling. Cell Physiol Biochem. 2017;44(2):423–35.

97. Polanco JC, Li C, Bodea LG, Martinez-Marmol R, Meunier FA, Gotz J. Amyloid-beta and tau complexity - towards improved biomarkers and targeted therapies. Nat Rev Neurol. 2018;14(1):22–39.

98. Liu J, Farmer JD, Jr., Lane WS, Friedman J, Weissman I, Schreiber SL. Calcineurin is a common target of cyclophilin-cyclosporin A and FKBP-FK506 complexes. Cell. 1991;66(4):807–15.

99. Wang Q, Wang J. Injection of bradykinin or cyclosporine A to hippocampus induces Alzheimer-like phosphorylation of Tau and abnormal behavior in rats. Chin Med J (Engl). 2002;115(6):884–7.

100. Yu DY, Luo J, Bu F, Song GJ, Zhang LQ, Wei Q. Inhibition of calcineurin by infusion of CsA causes hyperphosphorylation of tau and is accompanied by abnormal behavior in mice. Biol Chem. 2006;387(7):977–83.

101. Gong CX, Lidsky T, Wegiel J, Zuck L, Grundke-Iqbal I, Iqbal K. Phosphorylation of microtubule-associated protein tau is regulated by protein phosphatase 2A in mammalian brain. Implications for neurofibrillary degeneration in Alzheimer’s disease. J Biol Chem. 2000;275(8):5535–44.

102. Bialojan C, Takai A. Inhibitory effect of a marine-sponge toxin, okadaic acid, on protein phosphatases. Specificity and kinetics. Biochem J. 1988;256(1):283–90.

103. Swingle M, Ni L, Honkanen RE. Small-molecule inhibitors of ser/thr protein phosphatases: specificity, use and common forms of abuse. Methods Mol Biol. 2007;365:23–38.

104. Amundsen R, Asberg A, Ohm IK, Christensen H. Cyclosporine A- and tacrolimus-mediated inhibition of CYP3A4 and CYP3A5 in vitro. Drug Metab Dispos. 2012;40(4):655–61.

105. Serkova N, Brand A, Christians U, Leibfritz D. Evaluation of the effects of immunosuppressants on neuronal and glial cells in vitro by multinuclear magnetic resonance spectroscopy. Biochim Biophys Acta. 1996;1314(1-2):93–104.

106. Zhang F, Phiel CJ, Spece L, Gurvich N, Klein PS. Inhibitory phosphorylation of glycogen synthase kinase-3 (GSK-3) in response to lithium. Evidence for autoregulation of GSK-3. J Biol Chem. 2003;278(35):33067–77.

107. Ryves WJ, Harwood AJ. Lithium inhibits glycogen synthase kinase-3 by competition for magnesium. Biochem Biophys Res Commun. 2001;280(3):720–5.

108. Kase H, Iwahashi K, Nakanishi S, Matsuda Y, Yamada K, Takahashi M, et al. K-252 compounds, novel and potent inhibitors of protein kinase C and cyclic nucleotide-dependent protein kinases. Biochem Biophys Res Commun. 1987;142(2):436–40.

109. Thuret G, Chiquet C, Herrag S, Dumollard JM, Boudard D, Bednarz J, et al. Mechanisms of staurosporine induced apoptosis in a human corneal endothelial cell line. Br J Ophthalmol. 2003;87(3):346–52.

110. Green TP, Fennell M, Whittaker R, Curwen J, Jacobs V, Allen J, et al. Preclinical anticancer activity of the potent, oral Src inhibitor AZD0530. Molecular oncology. 2009;3(3):248–61.

111. Rubenstein R, Sharma DR, Chang B, Oumata N, Cam M, Vaucelle L, et al. Novel Mouse Tauopathy Model for Repetitive Mild Traumatic Brain Injury: Evaluation of Long-Term Effects on Cognition and Biomarker Levels After Therapeutic Inhibition of Tau Phosphorylation. Front Neurol. 2019;10:124.

112. Zhao Z, Wang L, Volk AG, Birch NW, Stoltz KL, Bartom ET, et al. Regulation of MLL/COMPASS stability through its proteolytic cleavage by taspase1 as a possible approach for clinical therapy of leukemia. Genes Dev. 2019;33(1-2):61–74.

113. Segura-Egea JJ, Jimenez-Rubio A, Rios-Santos JV, Velasco-Ortega E, Calvo-Gutierrez JR. In vitro inhibitory effect of EGTA on macrophage adhesion: endodontic implications. Journal of endodontics. 2003;29(3):211–3.

114. Knaryan VH, Samantaray S, Park S, Azuma M, Inoue J, Banik NL. SNJ-1945, a calpain inhibitor, protects SH-SY5Y cells against MPP(+) and rotenone. J Neurochem. 2014;130(2):280–90.

115. Henzing AJ, Dodson H, Reid JM, Kaufmann SH, Baxter RL, Earnshaw WC. Synthesis of novel caspase inhibitors for characterization of the active caspase proteome in vitro and in vivo. J Med Chem. 2006;49(26):7636–45.

116. Chopra P, Gupta S, Dastidar SG, Ray A. Development of cell death-based method for the selectivity screening of caspase-1 inhibitors. Cytotechnology. 2009;60(1-3):77.

